# Phosphoinositide-specific Phospholipase C 2 (SlPLC2) Facilitates Vesicle Formation and Modulates Immune Signaling in Tomato *Phytophthora infestans* Interactions

**DOI:** 10.1101/2025.09.21.677499

**Authors:** Enzo A. Perk, Juan Martín D’Ambrosio, Ignacio Cerrudo, Luciana Robuschi, Marcelo Juárez, Pilar Vélez, Verónica Mary, Martín Theumer, María Eugenia Segretin, Ana M. Laxalt

## Abstract

Phosphoinositide-specific phospholipase C (PI-PLC) is a signaling enzyme that hydrolyzes membrane phosphoinositide to generate lipid- and lipid-derived second messengers. In plants, PI-PLCs have been implicated in various physiological processes, including immunity. Tomato SlPLC2 was previously shown to be implicated in susceptibility to the necrotrophic fungus *Botrytis cinerea*. However, whether this observation extends to pathogens with different lifestyles and evolutionary origins, remains unknown. Here, we investigated SlPLC2 function during infection with *Phytophthora infestans*, a globally devastating oomycete and causal agent of potato and tomato late blight. CRISPR-Cas9 knockout *SlPLC2* plants showed significantly reduced disease symptoms, lower pathogen biomass, and decreased sporangia production, indicating impaired colonization. Upon infection, *SlPLC2* knockout lines displayed attenuated expression of salicylic acid (SA)- and jasmonic acid (JA)-responsive genes suggesting disrupted hormone signaling. Consistent with PLC role during plant defense, early immune responses such as hydrogen peroxide accumulation and callose deposition were reduced in knockout plants. At the cellular level, these plants allow fewer infection vesicles formation by *P. infestans*, accompanied by an increase in expression of a biotrophy-associated effector gene *PiAvrblb2*, suggesting impaired establishment of infection. In *Nicotiana benthamiana* SlPLC2-GFP localizes predominantly at the plasma membrane. Upon *P. infestans* inoculations, SlPLC2-GFP localizes to the membrane surrounding infection vesicles, where we also detected phosphoinositides PI4P and PI(4,5)P₂. Overexpression of *SlPLC2* in *Nicotiana benthamiana* enhanced susceptibility to *P. infestans*, reinforcing its positive role in colonization. Together, these findings identify SlPLC2 as a susceptibility factor associated with enhanced *P. infestans* colonization.

**Statement:** This study identifies SlPLC2 as a key susceptibility factor that supports *Phytophthora infestans* infection by coordinating vesicle formation and early immune responses in tomato.

## Introduction

Late blight, caused by the oomycete *Phytophthora infestans*, continues to inflict heavy economic losses on tomato (*Solanum lycopersicum*) and potato (*S. tuberosum*) despite decades of resistance breeding (Kamoun et al., 2006; Leesutthiphonchai et al., 2018). The pathogen disseminates in papillate sporangia that either germinate directly or release motile zoospores that encyst and germinate to produce germ-tubes (Judelson & Blanco, 2005; Judelson & Ah Fong, 2019). Germ-tubes form small appressoria beside trichomes or anticlinal cell walls (Avrova et al., 2008). Penetration generates a bulbous infection vesicle inside the initially invaded epidermal cell, from which intercellular hyphae emerges and spread into adjacent living cells (Boevink et al., 2020). In each newly colonized cell, digit-like haustoria differentiates, allowing the pathogen to maintain a biotrophic lifestyle (Boevink et al., 2020). These specialized infection structures are enveloped by a newly synthesized membrane that defines a unique host-pathogen interface. In the case of haustoria, this membrane is referred to as the extrahaustorial membrane (EHM) (Bozkurt et al., 2011). For infection vesicles, however, no specific name has been assigned, therefore, we will refer to it as the membrane surrounding the infection vesicle stage. Through these interfaces, *P. infestans* delivers an arsenal of effectors, of particularly interest are RXLR effectors-secreted proteins named after a conserved Arg-X-Leu-Arg motif that suppress immune responses and reprogram host cellular processes to favor colonization (Bozkurt et al., 2011; Dagdas et al., 2016; Avrova et al., 2008). The EHM displays a lipid and protein signature markedly different from the plasma membrane (PM) due to selective vesicle traffic that removes many PM constituents (Bozkurt and Kamoun 2020). EHM contains the phosphoinositide PI4P and PI(4,5)P₂, that are related to shaping the surface charge, curvature and membrane architecture (Shimada et al., 2019; Qin et al., 2020; Rausche et al., 2021). This tailored membrane landscape dictates the recruitment of specific host factors among which, remorin nanodomain scaffolds (Bozkurt et al., 2014; Turnbull et al., 2017), the late-endosomal RabG3c GTPase (Bozkurt et al., 2015), and the PI(4,5)P₂-specific 5-phosphatase StIPP that locally remodels phosphoinositide pools (Rausche et al., 2021). During the biotrophic phase, the pathogen keeps host cells alive suppressing immunity by delivering effectors across the EHM (Schornack et al., 2010; Whisson et al., 2016). A transcriptional re-programming of the pathogen, known as biotrophic-necrotrophic switch, trigger a secretion of toxins and microbial proteins that targets dicot plasma membranes, which results in necrotrophy, characterized by host-cell death (Wang et al., 2023). Finally, sporangiophores burst through stomata and lenticels, releasing new sporangia for dispersion (Judelson & Ah Fong, 2019). Yet, even after state-of-the-art ultrastructural, lipidomic and proteomic surveys of EHMs and related host-derived interfaces (Caillaud et al., 2014; Bozkurt & Kamoun, 2020), the host lipid-modifying enzymes and trafficking pathways that sculpt their distinctive membrane landscape remain largely unidentified.

One of the routes by which plants convert phosphoinositide signals into immune responses involves the hydrolysis of PI4P and PI(4,5)P₂ by phosphoinositide-specific phospholipase C (PLC) (Laxalt et al., 2025). Activated PLC hydrolyses either PI4P or PI(4,5)P₂ to produce diacylglycerol (DAG) and soluble inositol phosphates (IP₂/IP₃). These compounds mobilize vacuolar Ca²⁺ stores, contributing to the generation of calcium signatures characteristic of pattern-triggered immunity. In parallel, DAG is rapidly phosphorylated by DAG kinase (DGK) to form phosphatidic acid (PA). In Arabidopsis, upon flg22 perception, PLC2 and DGK5 are required for oxidative burst (D’Ambrosio et al., 2017; Kong et al., 2024). The PLC/DGK-derived PA stabilizes and activates the NADPH oxidase RBOHD at the plasma membrane to enhance ROS production (Zhang et al., 2009; Qi et al., 2024 and Kong et al., 2024). Through this cascade, PLC2 and DGK5 link phosphoinositide metabolism to the production of Ca²⁺ and ROS signals that are involved in plant immunity. The conversion of PI4P- and PI(4,5)P₂ into PA provides a mechanism to dynamically reprogram the electrostatic and signaling properties of the plasma membrane during immune activation. Based on our observations in tomato, we propose that PLCs may contribute to local membrane remodeling by depleting anionic phosphoinositides and generating second messengers that regulate Ca²⁺ influx, ROS production, and vesicle dynamics during immune responses.

Tomato encodes six PLC genes (*SlPLC1-SlPLC6*). Loss-of-function of *SlPLC2*-via either virus-induced gene silencing or CRISPR/Cas9 editing resulted in plants less susceptible to *Botrytis cinerea* (Gonorazky et al., 2016; Perk et al., 2023). Thus, SlPLC2 was proposed as a susceptibility (S) gene (Perk et al., 2023). A S gene is required by the pathogen for a successful colonization, so that recessive loss-of-function alleles confer broad and often durable resistance (van Schie & Takken, 2014; Pavan et al., 2010). In a similar scenario, we hypothesize that SlPLCs could facilitates *P. infestans* colonization. We show that *SlPLC2* is up-regulated during infection, promotes infection-vesicle formation and pathogen proliferation, modulates ROS production and SA/JA gene expression. SlPLC2 localizes in infection vesicles where PI4P- and PI(4,5)P₂ are present. These findings position *SlPLC2* as a susceptibility factor that orchestrates immune signaling and lipid dynamics at host-pathogen interfaces and thus represents a promising new target for durable late-blight resistance.

## Results

### *SlPLC* transcripts are differentially induced during *P. infestans* infection

To investigate whether the *SlPLC* gene family participates in the tomato immune response to *P. infestans*, we analyzed their expression dynamics following pathogen inoculation. This analysis builds upon previous findings showing that *SlPLC* genes are transcriptionally activated in response to the necrotrophic pathogen *B. cinerea* (Gonorazky et al., 2016). To assess whether phosphoinositide signaling is broadly engaged during pathogen attack, we examined the transcriptional dynamics of the *SlPLC* gene family during infection by *P. infestans*. We performed time-course RT-qPCR assays using RNA extracted from whole leaflets drop-inoculated with a zoospore suspension placed on either side of the central vein. Transcript levels of *SlPLC1-6* were measured at 1, 2, 4, and 6 days post-inoculation (dpi) and compared to non-infected controls. As shown in Supplemental Figure S1, no major transcriptional changes were detected between 1 and 4 dpi, indicating that the early stages of infection do not substantially affect *SlPLC* transcript abundance at the whole-leaf level. A clear induction of specific *SlPLC* genes was only observed at 6 dpi, coinciding with the radial expansion of lesions and widespread pathogen colonization. This late transcriptional response likely reflects the spatial heterogeneity of infection foci within the leaflet: while some regions are undergoing necrosis, others are being newly colonized and still experience biotrophic invasion. As a result, the observed expression changes at 6 dpi integrate signals from diverse infection stages across the tissue. By 6 dpi, three members of the family—*SlPLC1*, *SlPLC2*, and *SlPLC3*—exhibited significantly elevated transcript levels (Figure 1). Signaling enzymes often display a property where treatments that activate them lead to an enhancement of their gene expression. This response is thought to be a positive feedback mechanism that primes the cell for further stimulation (Hirt, 1999; Yamamoto,1998). Among the *SlPLC* gene family members analyzed, *SlPLC2* emerged as a candidate since it was previously reported as a susceptibility gene during *B. cinerea* infections (Gonorazky et al., 2016; Perk et al., 2023) and is transcriptionally induced during *P. infestans* infection (Figure 1).

**Figure 1.**
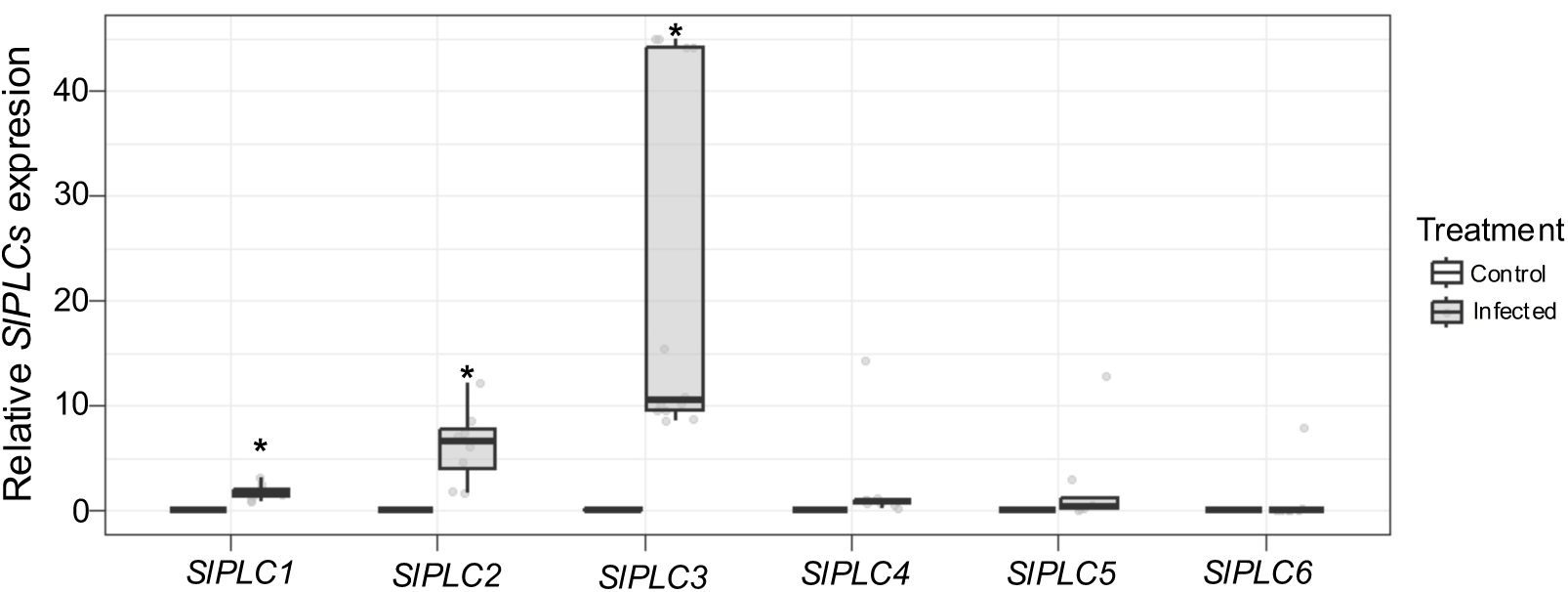
*SlPLC1*, *SlPLC2*, and *SlPLC3* transcript levels are induced during *P. infestans* infection. Tomato leaflets from 6-week-old plants were inoculated with *P. infestans* (applied as two 10 µL droplets of 2×10^5^ zoospores per mL solution on either side of the central vein) or non-infected. Samples were collected at 6 dpi for RNA extraction and gene expression analysis. Transcript levels of *SlPLC1* to *SlPLC6* were measured by RT-qPCR and normalized to *SlActin*. Data represent mean ± standard error (SE) of eight biological replicates for infected samples and four for non-infected plants. Asterisks indicate statistically significant differences compared to non-infected plants (Dunnett’s post-hoc test, * = P < 0.05).

### *SlPLC2*-knock out tomato plants are less susceptible to *P. infestans*

To assess SlPLC2 functional contribution during *P. infestans* infection, we analyzed two independent *SlPLC2* knockout lines (KO-6 and KO-17), generated using CRISPR-Cas9 technology reported earlier (Perk et al., 2023). Six-week-old tomato plants were drop-inoculated with *P. infestans* zoospores and disease symptoms were evaluated at 6 dpi. Both KO lines consistently developed significantly smaller necrotic lesions compared to WT plants (Figure 2 A, B), indicating reduced disease susceptibility. To validate this phenotype at the molecular level, we measured *P. infestans* biomass via RT-qPCR targeting the pathogen reference gene *PiEF1-α*. KO lines showed significantly lower pathogen transcript levels relative to WT plants (Fig. 2 C). As expected, *SlPLC2* transcripts were undetectable in KO lines due to nonsense-mediated decay confirming the effectiveness of the gene knockout (Supplemental Figure 1). The expression levels of the remaining *SlPLC* genes were comparable to those in the WT, suggesting that no compensatory mechanisms were activated (Supplemental Figure S1).

**Figure 2.**
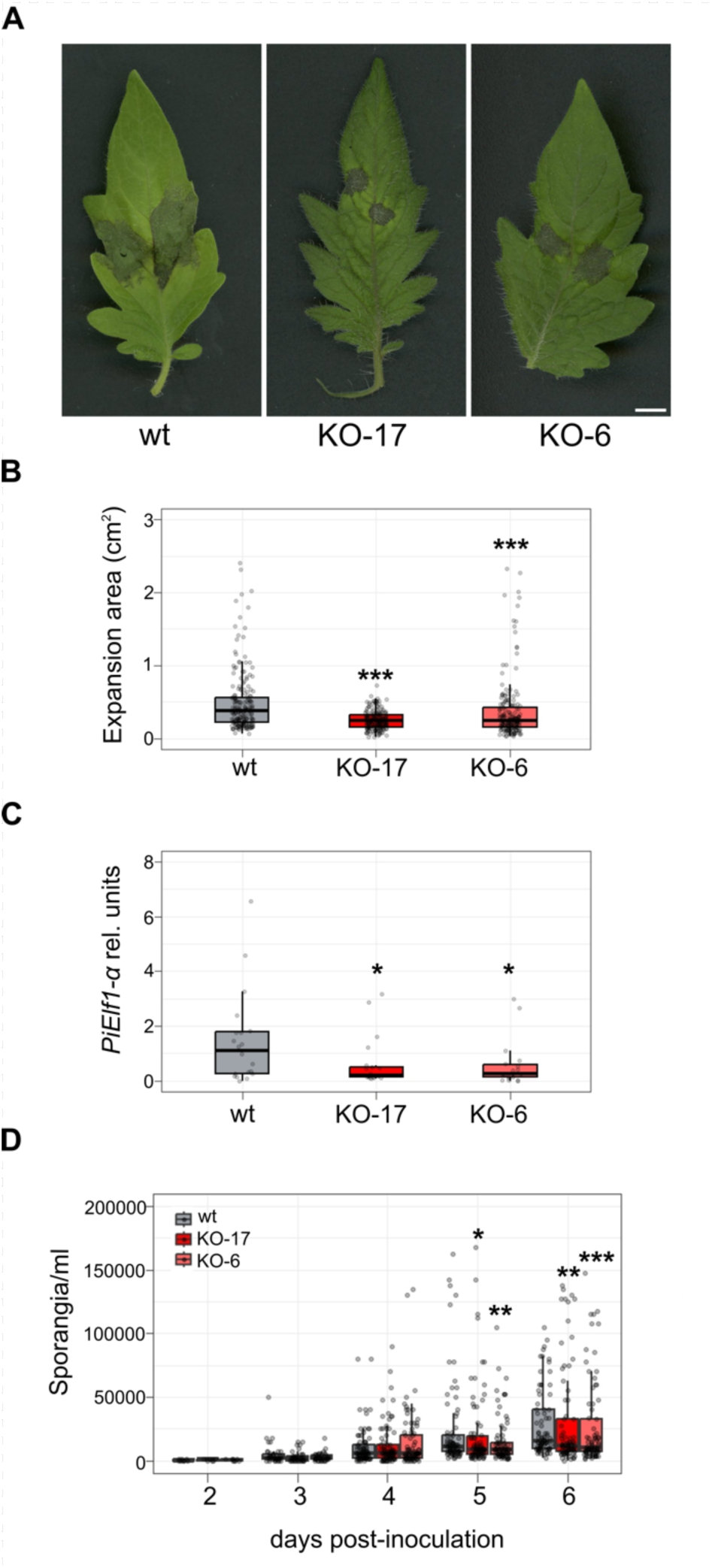
*SlPLC2* knockout lines display reduced susceptibility to *P. infestans*. Detach leaves of WT and *SlPLC2* knockout lines (KO-6 and KO-17) were inoculated on the abaxial side with *P. infestans* zoospores (a 10 µL droplet containing 2 × 10⁵ zoospores per mL) on either side of the central vein. Disease progression was assessed at 6 dpi by comparing lesion development and quantifying pathogen biomass. **(A)** Representative images of infected leaflets at 6 dpi. KO lines developed visibly smaller necrotic lesions compared to wt. Scale bar = 0.5 cm. **(B)** Lesion area was quantified from images in (A) using ImageJ. Data represent the mean ± SE of 300-400 inoculation sites pooled from three independent experiments. Asterisks indicate significant differences compared to WT (Dunnett’s post-hoc test, *** = *P* < 0.0005). **(C)** Pathogen biomass was determined by RT-qPCR amplification of the *P. infestans* elongation factor gene (*PiEF1-α*), normalized against tomato *SlACTIN* expression from infected and mock-treated leaflets at 6 dpi (n = 12 infected, n = 8 mock), Asterisks indicate statistical significance from WT plants (Dunnett’s post-hoc test, * = P < 0.05). (D) Time-course of sporulation in leaf discs from WT and SlPLC2 knockout lines (KO-6 and KO-17) following *P. infestans* inoculation. Leaf discs were drop-inoculated with 200 zoospores in 10 µL droplets. Sporangia were harvested and quantified at the indicated timepoints. Data are presented as box plots representing three independent experiments. Statistical comparisons at 5 dpi and 6 dpi were performed using the Kruskal-Wallis test followed by pairwise Wilcoxon tests. Asterisks denote statistically significant differences (* = p ≦ 0.05; ** = p ≦ 0.01; *** = p ≦ 0.001) based on the Wilcoxon test.

Then, we evaluated the formation of reproductive structures at later stages of infection. As such, sporulation is not only a marker of pathogen fitness but also reflects the overall progression and success of infection. To evaluate if *SlPLC2* KO plants impair *P. infestans* reproduction, we quantified sporangia production at 2,3,4,5 and 6 dpi. As shown in Fig. 2 D, KO lines exhibited a significant reduction in sporangia quantification relative to WT. The reduced sporangia production in SlPLC2 KO lines indicates that pathogen development is impaired, although the precise stage at which this occurs remains to be determined.

### Attenuated hormonal responses to *P. infestans* in *SlPLC2* knockout tomato plants

To explore the mechanisms underlying the reduced susceptibility observed in *SlPLC2* KO plants, we first characterized these responses in WT plants by monitoring the transcriptional dynamics of hormone-responsive gene markers. Salicylic acid (SA) and jasmonic acid (JA) play central, yet often antagonistic, roles in coordinating plant immunity. SA is typically associated with resistance to biotrophic and hemibiotrophic pathogens, while JA predominantly regulates responses against necrotrophs and insect herbivores (Glazebrook, 2005; Mur et al., 2006; Robert-Seilaniantz et al., 2011). However, the crosstalk between these pathways is dynamic and context-dependent, especially during hemibiotrophic infections such as those caused by *P. infestans*, where hormonal regulation is still not fully understood (Halim et al., 2009; Spoel & Dong, 2008). For the SA pathway, the defense marker gene *SlPR1a* was significantly induced at 2 dpi, with transcript levels continuing to rise by 6 dpi (Supplementary Figure S2A, Figure 3A). In parallel, expression of the SA biosynthetic gene *SlPAL* also increased over the course of infection (Figure 3B). However, free SA levels did not increase at 6 dpi (Figure 3C), suggesting that transcriptional activation of the SA pathway does not necessarily translate into detectable changes in free hormone accumulation at this stage. This discrepancy could reflect tight post-transcriptional or metabolic control of SA levels, or the rapid conversion of SA into conjugated or inactive forms not detected in our measurements.

**Figure 3.**
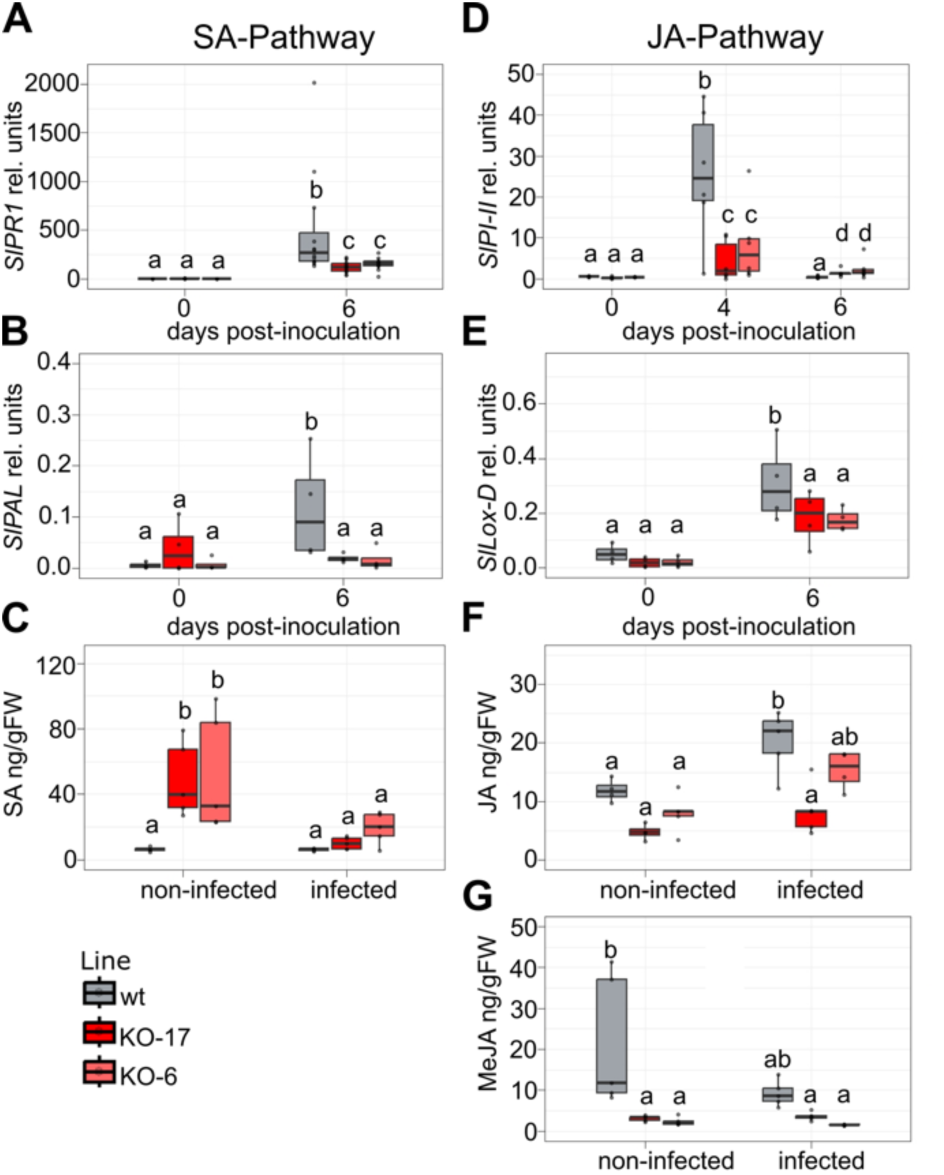
Hormonal imbalance and altered transcriptional responses in *SlPLC2* knockout lines during *P. infestans* infection. WT and *SlPLC2* knockout tomato lines (KO-6 and KO-17) were inoculated with *P. infestans* zoospores (2 × 10^3^ per leaflet, applied as two 10 µL droplets from a 2×10^5^ solution on either side of the central vein). **(A-B-D-E)** Transcriptional responses were analyzed by RT-qPCR at different dpi. Values were normalized to *SlACTIN*. **(A)** *SlPR1a* (salicylic acid marker), **(B)** *SlPAL* (SA biosynthesis), **(D)** *SlPI-II* (jasmonic acid marker), **(E)** *SlLOXD* (JA biosynthesis). Data represent the mean ± SE of biological replicates (n = 12 for infected samples at 6 dpi, n = 8 at 4 dpi, n = 4 for mock controls). Different letters indicate statistical differences (Dunnett’s test, * = P < 0.05). **(C-F-G)** Quantification of endogenous hormone levels by LC-MS/MS in aerial tissues at 6 dpi in infected or non-infected plants. Results were expressed as nanograms per milligram of fresh weight. **(C)** Salicylic acid (SA), **(F)** Jasmonic acid (JA), **(G)** Methyl jasmonate (MeJA). Data represent the mean ± SE of biological replicates (n = 5). Different letters indicate statistical differences (Dunnett’s test, * = P < 0.05).

We then analyzed JA-related responses. The JA-responsive gene *SlPI-II* remained low during early stages of infection, became strongly induced at 4 dpi, and returned to basal levels by 6 dpi (Supplementary Figure S2 B). This transient induction is consistent with a short-lived activation of JA signaling, likely associated with the onset of the necrotrophic phase and the presence of damage-associated molecular patterns (DAMPs). Upregulation of the JA biosynthetic gene *SlLoxD* (Figure 3E), together with a significant increase in JA levels at 6 dpi in WT plants (Figure 3F), indicates activation of the JA pathway during the later stages of infection. In contrast, levels of methyl jasmonate (MeJA) did not show significant variation at this stage (Figure 3G).

To determine whether hormonal responses were altered in the absence of SlPLC2, we examined the expression of hormone-responsive genes, biosynthetic markers, and hormone levels in KO lines. In the SA pathway, the expression of the SA-responsive gene *SlPR1a* was significantly reduced in KO plants compared to WT at 6 dpi (Figure 3A). This attenuated induction may reflect an impaired in SA signal transduction downstream of hormone perception. Similarly, the expression of the SA biosynthetic gene *SlPAL* was also lower in KO lines relative to WT (Figure 3B). Despite these transcriptional differences, total SA levels remained comparable between genotypes at 6 dpi (Figure 3C). To our surprise KO non-infected detached leaves after six days exhibited significantly higher SA levels than WT (Figure 3C).

In the JA pathway, the expression of the JA-responsive gene *SlPI-II* was significantly lower in KO plants compared to WT at 4 dpi (Figure 3D) and never reached WT levels throughout the time course. Interestingly, while transcript levels declined rapidly in WT by 6 dpi, KO lines maintained slightly higher levels at this later stage. This suggests a slower decline in *SlPI-II* expression rather than sustained activation. Such a profile may reflect delayed resolution of the response in the absence of SlPLC2. Consistently, the expression of the JA biosynthetic gene *SlLoxD* was also lower in KO plants (Figure 3E), JA levels during infection were significantly reduced in one line and statistically intermediate in the other, relative to WT (Figure 3F). These results show that JA pathway activation is reduced in KO plants during infection. KO plants exhibited significantly reduced levels of methyl jasmonate (MeJA) under non-infected conditions, which correlates with elevated SA levels (Figure 3G).

WT plants and both *SlPLC2* knockout lines exhibited comparable basal levels of SA, JA and MeJA (Supplementary Figure 3). Hormone quantification was performed using detached leaves that were immediately processed after excision to minimize artifactual changes in endogenous hormone levels.

To assess whether the hormonal alterations observed in KO plants were specific to defense-related pathways, we also measured the levels of other phytohormones such as abscisic acid (ABA) and indole-3-acetic acid (IAA). Their levels remained unchanged in KO compared to WT plants at 6 dpi (Supplementary Figure S 4 A B), suggesting that the effects of SlPLC2 disruption are specific to SA and JA pathways rather than reflecting a broader hormonal imbalance. Altogether, these findings indicate that absence of SlPLC2 leads to altered SA- and JA-related responses.

### ROS production and callose deposition are impaired in *SlPLC2* KO plants

Given the reduced susceptibility observed in *SlPLC2* KO plants and altered hormonal balance and gene expression, we investigated whether this phenotype is associated with alterations in immune signaling events regulated by SlPLC2. PI-PLCs are required for ROS production via NADPH oxidase RBOHD during pathogen perception (Laxalt et al., 2025). For that, we assessed ROS accumulation at 1 dpi, through 3,3’-diaminobenzidine (DAB) staining. While WT plants displayed strong hydrogen peroxide (H₂O₂) accumulation around infection sites KO lines exhibited significantly reduced DAB staining (Figure 4 A B), consistent with a disruption in oxidative signaling due to the absence of SlPLC2.

**Figure 4.**
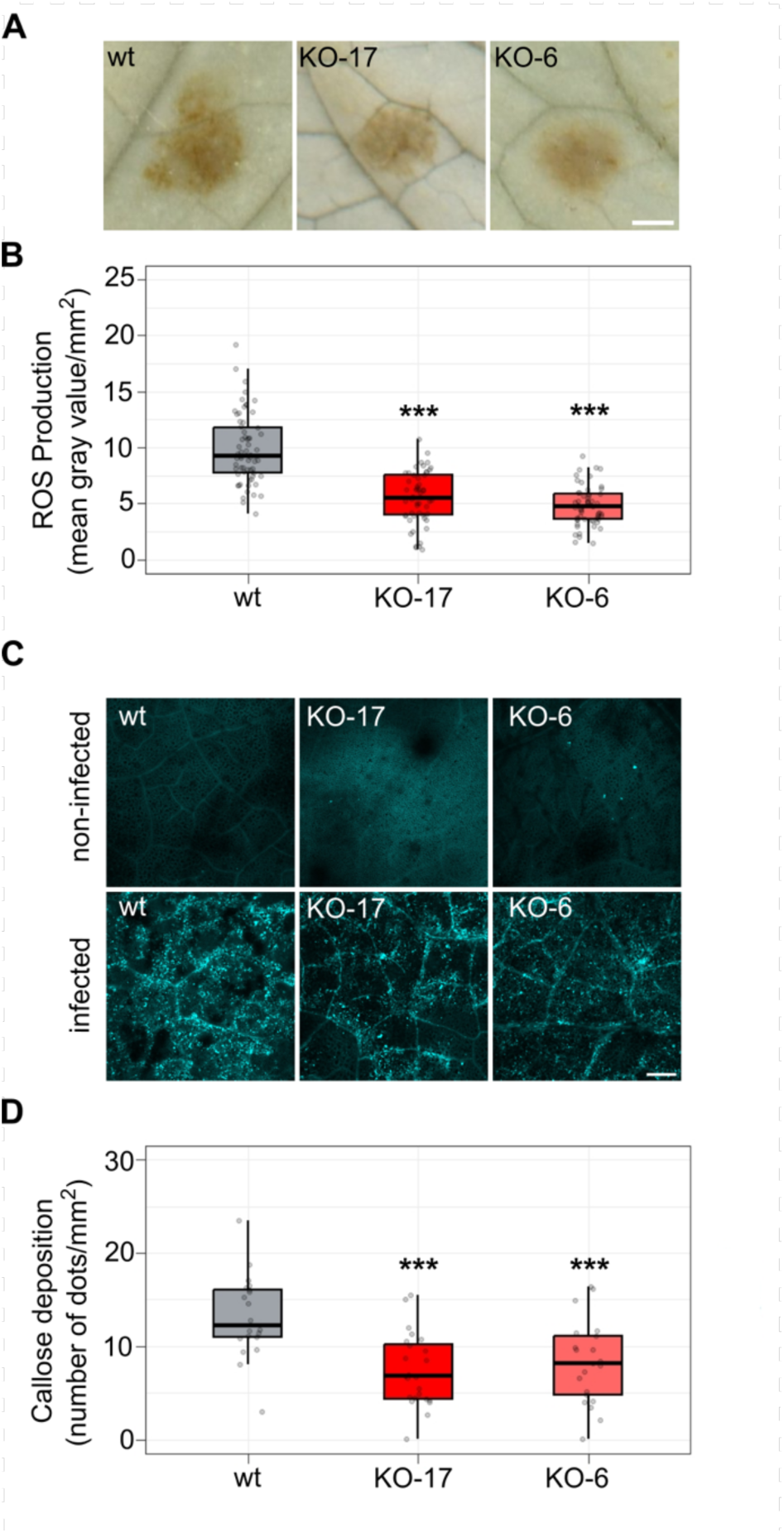
Attenuated ROS production and callose deposition in *P. infestans* inoculated *SlPLC2* knockout plants. Leaflets from wild-type (WT) and *SlPLC2* knockout tomato lines (KO-6 and KO-17) were inoculated with *P. infestans* zoospores (2 × 10⁵ zoospores per mL as a 10 µL droplet). **(A)** Detection of hydrogen peroxide (H₂O₂), 1 dpi using 3,3′-diaminobenzidine (DAB). A representative image of H₂O₂ accumulation, (visualized as brown precipitate). Scale bar = 4 mm. **(B)** Quantification of DAB staining intensity using ImageJ, expressed as mean gray value per mm². **(C)** Detection of callose in 2 dpi infected leaves by aniline blue staining under epifluorescence microscopy (excitation 365/10 nm; emission 460/50 nm). A representative image of callose deposition is shown. Scale bar = 200 µm. **(D)** Quantification of callose deposition was carried out using ImageJ by measuring the number of fluorescent callose deposits per mm². In all panels, data represent mean ± SE of at least three independent biological replicates. Asterisks indicate statistically significant differences compared to WT (Dunnett’s test, *P* < 0.05).

One of the key defense processes regulated downstream of ROS is callose deposition (Luna et al., 2011; Kadota et al., 2014). We assessed whether this defense mechanism was also impaired in KO lines. Leaves were stained with aniline blue at 2 dpi to visualize callose deposition. While WT plants exhibited robust callose accumulation around infection sites, KO lines showed a marked reduction in both signal intensity and coverage (Figure 4 C D), indicating that ROS-dependent structural defenses are also attenuated in the absence of SlPLC2. Together, these findings suggest that SlPLC2 contributes to immune signaling, influencing both ROS accumulation and the activation of downstream defense pathways, including hormonal responses and callose deposition.

### Delayed progression of *P. infestans* infection in *SlPLC2* knockout tomato plants

To characterize the initial stage of *P. infestans* colonization in tomato, we focused on the formation of infection vesicles—specialized intracellular structures that represent the first interface between the pathogen and the host cell. To enable microscopic visualization of these early infection events, we used the *P. infestans* 88069tD strain, which expresses tdTomato (Red channel) allowing direct observation of oomycete structures within plant tissue (Figure 5 A B). Quantification of infection vesicles on tomato leaves revealed a significant reduction in the number of infection vesicles formed in the KO lines compared to WT plants (Figure 5 C). This difference suggests that SlPLC2 may contribute to the infection vesicle formation.

**Figure 5.**
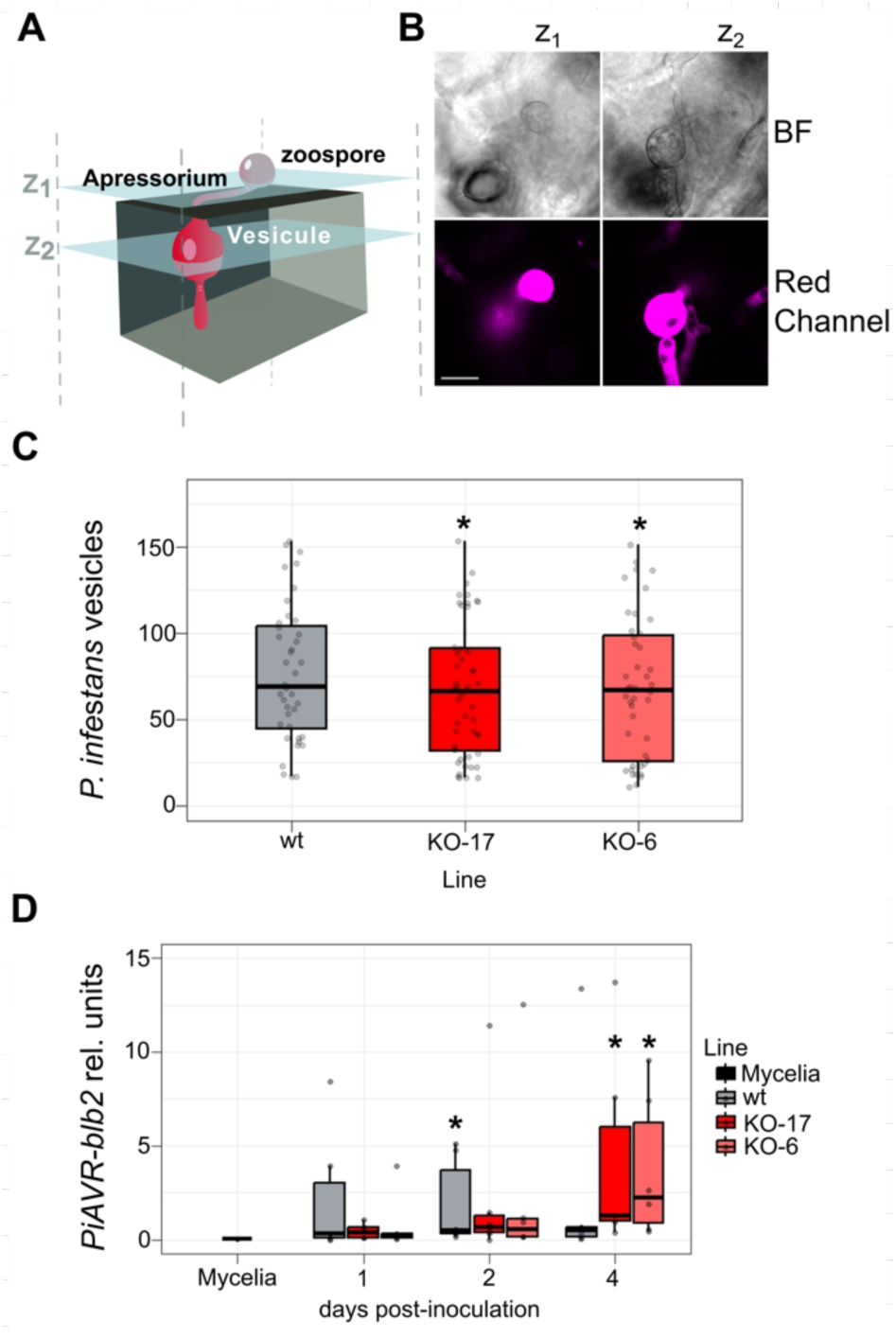
Altered infection dynamics of *P. infestans* in *SlLC2 KO* Tomato Lines. Detached leaflets from WT and KO tomato plants were drop-inoculated on the abaxial surface with 10 µL of a suspension containing 2 × 10^3^ zoospores of *P. infestans PinSL-19* or 88069td expressing a cytoplasmic tdTomato (RFP). Infected leaves were incubated under high humidity conditions and sampled at 1-, 2-, and 4-days post-inoculation (dpi). Then leaf 1 cm diameter disk were exscinded from the infection site to further analysis. **(A)** Schematic representation of the *P. infestans* infection process showing an epidermal cell with a germinating zoospore forming an appressorium, a spherical infection vesicle, and a primary hypha. Z1 and Z2 correspond to different focal planes from the microscopy image. **(B)** Representative confocal images of an infection vesicle at 1 dpi in *S. lycopersicum* leaf tissue inoculated with *P. infestans* strain 88069td expressing tdTomato. Optical sections are shown at Z1 and Z2 planes. Images were acquired with a Nikon C1siR laser-scanning confocal microscope equipped with a Plan Fluor 40× objective and controlled with Nikon EZ-C1 software. tdTomato fluorescence was detected in the RFP channel (excitation 543 nm; emission 560-615 nm). Laser intensity, detector gain, and offset were kept constant across samples within the experiment. Scale bar = 10 µm. **(C)** Quantification of infection vesicles of 88069td at 1 dpi in WT and KO genotypes in tomato. Infection vesicles were visualized by epifluorescence microscopy, and the number of vesicle-containing cells was quantified from at least three fields per disk (24 disks per biological replicate, three biological replicates). **(D)** Transcript levels of the *P. infestans* effector gene *PiAvrblb2*, measured by RT-qPCR at plate, 1, 2, and 4 dpi in infected leaf tissue. For expression analyses, tomato leaflets were infected with *PiNSL-19* strain and sampled at the indicated time points. From each leaflet, a 1 cm diameter disk was excised using a cork borer and immediately frozen in liquid nitrogen. For each biological replicate (n = 4), total RNA was extracted from a pool of 8 disks. *PiAvrblb2* transcript levels were quantified using gene-specific primers (Supplemental Table S1), normalized to the constitutive *P. infestans* gene *PiEF1-α.* The Mycelia (plate culture) condition was used as the control. For each line × time post-inoculation (dpi) combination, an independent comparison against Mycelia was performed using a two-sided Welch’s t-test (ttest_ind, equal_var=False), without any global model. To control the family-wise error rate across multiple comparisons, *p*-values were adjusted using the Holm-Bonferroni method (multipletests, method=“holm”). Statistical significance was set at α = 0.05 (adjusted).

To gain deeper insight into infection progression, we quantified PiAvrblb2, an RXLR-type effector typically expressed during the biotrophic phase of *P. infestans*. Expression values (relative to the endogenous pathogen reference PiELF-1) were directly compared against plate-grown mycelia (Mycelia) as the reference condition. At 1 dpi, PiAvrblb2 levels did not differ from Mycelia in any line. At 2 dpi, WT plants showed a significant induction vs Mycelia, whereas KO lines did not (Figure 5D). By 4 dpi, PiAvrblb2 in WT was no longer different from Mycelia, while both KO lines exhibited a significant induction (Figure 5D). This kinetics suggest that the onset of infection is delayed in KO backgrounds relative to WT. At 4 dpi, the constitutive pathogen transcript PiEF1-α accumulated in WT but remained lower in the KO lines, indicating delayed pathogen progression in the absence of SlPLC2 (Supplemental Figure S5). Together with the reduced infection-vesicle formation, lower pathogen load, decreased sporangia development, and delayed induction of biotrophy-associated effectors such as PiAvrblb2, these findings support a role for SlPLC2 in facilitating early host colonization by *P. infestans*. Interestingly, members of the PiAvrblb2 effector family have been shown to promote *P. infestans* virulence by targeting multiple layers of host immunity, including MAPK signaling, apoplastic protease activity, and Ca²⁺-dependent signaling; thus, the delayed induction of PiAvrblb2 in the KO background is likely to further compromise pathogen establishment (Bozkurt et al., 2011, Naveed et al., 2019, Lee et al., 2023).

### SlPLC2 localizes to infection vesicles and promotes host susceptibility

The observed alteration in infection establishment prompted us to investigate how SlPLC2 contributes to the first stages of *P. infestans* colonization at the cellular level. To do so, we investigated its subcellular localization by transiently expressing SlPLC2 fused to GFP in *N. benthamiana* leaves. Under non-infected conditions, SlPLC2-GFP fluorescence was predominantly localized at the plasma membrane, while free GFP exhibited diffuse cytosolic and nuclear distribution (Supplementary Figure S6A). A weaker but detectable SlPLC2-GFP signal was also observed in the cytosol, indicating that the fusion protein localizes to both membranes and the cytosol. Expression of the SlPLC2-GFP construct was validated by RT-PCR and western blotting confirming the presence of the expected transcript and tagged protein (Supplementary Figure S6 B C). Following inoculation with strain 88069td, at 1 dpi a subset of SlPLC2-GFP signal localized to discrete punctate structures that partially overlapped with RFP-labeled infection vesicles (Figure 6 A), indicating localization to interface membranes. In contrast, GFP alone was not observed at the sites of infection vesicles (Figure 6 A). These observations highlight the specific association of SlPLC2-GFP with the plasma membrane and its localization toward host-pathogen interface membranes at the site of infection vesicles.

**Figure 6.**
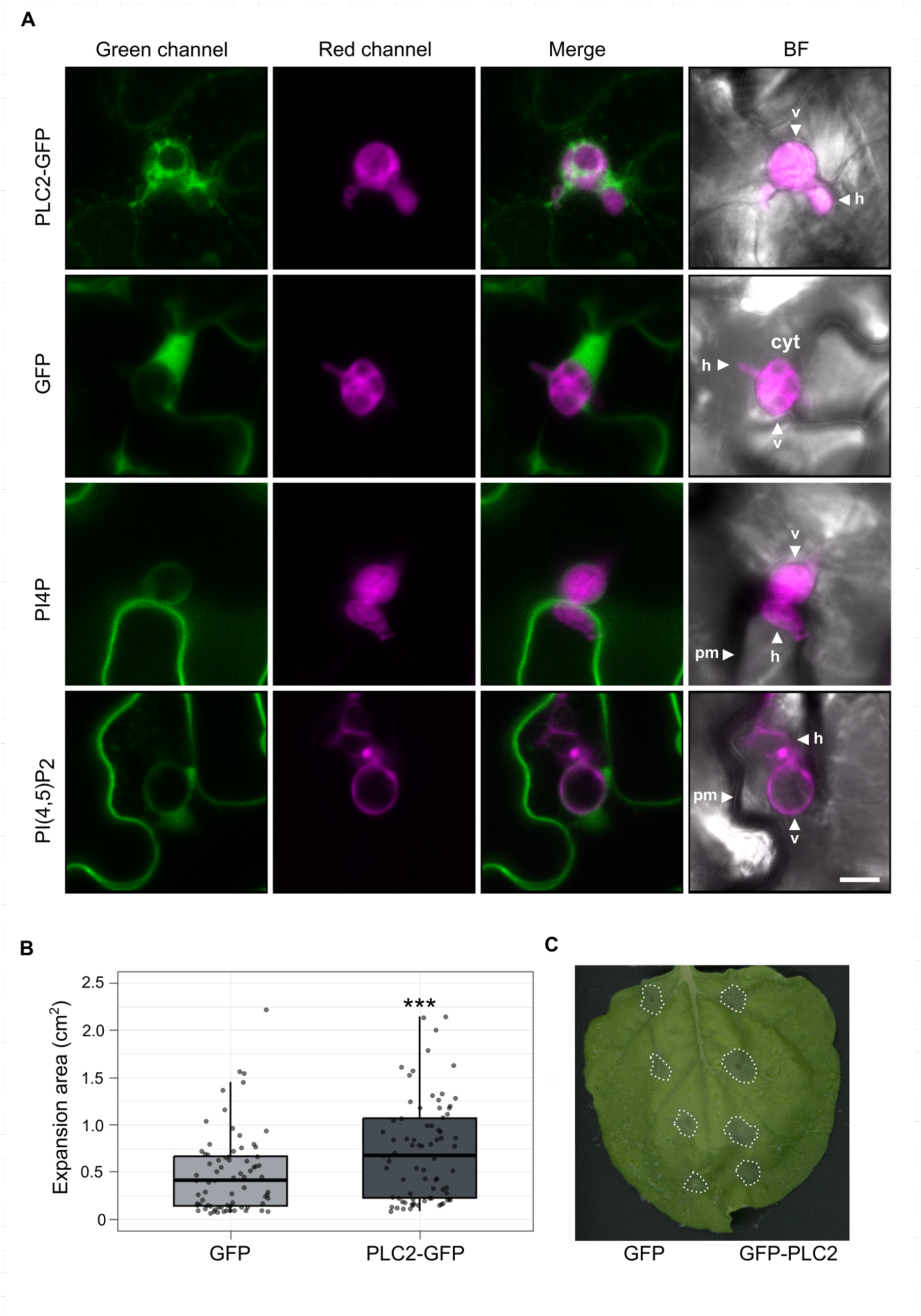
Subcellular localization of SlPLC2-GFP, PI4P and PI(4,5)P₂ in *P. infestans*-infection vesicles in *Nicotiana benthamiana*. Detached *Nicotiana benthamiana* leaves were agroinfiltrated with constructs expressing SlPLC2-GFP, GFP, the PI4P sensor (mCitrine-2xPH_FAPP1_), or the PI(4,5)P₂ sensor (mCitrine-2xPH_PLC_). At 48 hours post-agroinfiltration, leaves were drop-inoculated on the abaxial surface with 10 µL of a suspension containing 2 × 10⁵ zoospores per mL of *P. infestans*. Infected leaves were incubated under high humidity conditions and imaged by confocal microscopy at 1-day post-inoculation (dpi). **(A)** Confocal images of *N. benthamiana* epidermal cells at 1 dpi with *P. infestans* 88069td, showing the localization of SlPLC2-GFP, GFP, the PI4P sensor (mCitrine-2xPH(_FAPP1_) and the PI(4,5)P₂ sensor (mCitrine-2XPH_PLC_). SlPLC2-GFP accumulates in punctate structures partially overlapping with RFP-labeled infection vesicles. GFP alone localizes to the cytosol and nuclei. PI4P and PI(4,5)P₂ sensors show membrane-associated and cytosolic signals, including enrichment around infection vesicles Scale bar = 10 µm. cyt : Cytosol, pm : Plasma membrane, v : Infection vesicle, h : hyphae. **(B)** Quantification of lesion size at 6 dpi in *N. benthamiana* leaves transiently expressing SlPLC2-GFP or GFP infected with *P. infestans (PinSL-19)*. Infections were performed using the same inoculation method and zoospore concentration as in panel A. **(C)** Representative picture, Scale bar = 1 cm.

To further investigate whether the distribution of SlPLC2 correlates with changes in the spatial distribution of its potential substrates, we analyzed the localization of phosphoinositides-PI4P and PI(4,5)P₂-using fluorescent lipid biosensors, which consist of specific lipid-binding domains fused to a fluorescent protein (Simon et al., 2014) expressed in *N. benthamiana* leaves. The PI4P sensor was strongly localized at the plasma membrane under non-infected conditions (Supplementary Figure S6 D), but upon infection, it also localized at the membrane around infection vesicles (Figure 6 A). The PI(4,5)P₂ sensor showed a partial membrane and cytosolic distribution in both infected and non-infected cells and was detectable at the membrane surrounding infection vesicles (Figure 6 A, Supplementary Figure S6 D). Both sensors marked membranes associated with infection vesicles, but with reduced signal intensity relative to the plasma membrane, reflecting the distinct lipid composition of these newly formed host-pathogen interface membranes (Figure 6A, Supplementary Figure 7). SlPLC2 localizes the plasma membrane and to infection-associated vesicles during infection, whereas the GFP control remains confined to the cytosol (Supplementary Figure 7).

Finally, reasoning that *SlPLC2* KO lines are less susceptible to *P. infestans* infection, we hypothesized that overexpression of SlPLC2 might have the opposite effect. To test this, we assayed disease progression in *N. benthamiana* leaves transiently overexpressing SlPLC2-GFP or GFP alone. At 6 dpi, leaves expressing SlPLC2-GFP developed significantly larger necrotic lesions compared to GFP-expressing controls (Figure 6 B), indicating that elevated SlPLC2 expression can enhance disease susceptibility.

## Discussion

Our results identify *SlPLC2* as a susceptibility gene in tomato that modulates both the timing and magnitude of immune responses during *P. infestans* infection. Using CRISPR/Cas9-generated knockout lines in combination with gene expression and hormonal profiling, cellular imaging, and functional assays, we demonstrate that *SlPLC2*-KO plants are less susceptible to infection. This reduced susceptibility correlates with attenuated immune responses and impaired formation of infection vesicles. Notably, SlPLC2-GFP localizes to the plasma membrane in uninfected tissue but is recruited to infection vesicles during early colonization, suggesting that SlPLC2 operates at the host-pathogen interface. Together, these findings support a role for SlPLC2 in immune signal transduction and membrane remodeling during the establishment of infection. This expands previous work implicating phosphoinositide-modifying enzymes as central regulators of plant immunity (Abd-El-Haliem & Joosten, 2017; Rausche et al., 2021; Heilmann & Heilmann, 2022; Robuschi et al., 2024; Qin et al., 2020; Shimada et al., 2019, Guyon et al., 2025).

### Transcriptional induction of *SlPLC2* and related genes during infection

Gene expression analysis revealed that SlPLC2 is strongly induced at 6 dpi, coinciding with the appearance of necrotic lesions and sporulation. This transcriptional activation occurs alongside that of other PLC paralogs such as SlPLC1 and SlPLC3, suggesting a broader engagement of the PLC family during infection. SlPLC2 was prioritized for functional studies based on previous evidence linking it to susceptibility to *B. cinerea* (Gonorazky et al., 2016; Perk et al., 2023). In the context of *P. infestans*-a hemibiotroph with a radially progressing infection front-different leaflet zones likely correspond to biotrophic or necrotrophic stages. Therefore, SlPLC2 upregulation at 6 dpi may reflect its involvement across multiple infection phases. Notably, disruption of SlPLC2 compromises disease development in interactions with both *B. cinerea* and *P. infestans*, supporting its role as a conserved vulnerability factor targeted by distinct pathogen types. Together, these observations position SlPLC2 as a transcriptionally responsive susceptibility gene that contributes to infection progression across pathogen lifestyles, and phylogenetically unrelated.

### Hormonal reprogramming in *SlPLC2* knock-out plants

Hormonal profiling of *SlPLC2* KO plants during *P. infestans* infection revealed an altered immune signaling landscape. While total SA and MeJA levels did not differ consistently significantly between genotypes during infection, and JA levels were reduced in one of the two KO lines, the expression of key SA-associated and JA-associated genes was significantly lower in KO lines compared to WT. Given the well-characterized antagonism between SA and JA signaling (Glazebrook, 2005; Robert-Seilaniantz et al., 2011), the altered gene expression observed in KO plants likely contributes to enhanced resistance. However, another plausible explanation is that the reduced JA and SA signaling observed in *SlPLC2* KO plants could also reflect a diminished pathogen proliferation rather than a direct regulatory role of SlPLC2 in hormone signaling.

Further supporting a role for *SlPLC2* in modulating hormone-dependent defenses, several studies in tomato have shown that perturbations in SA and JA pathways can result in either increased susceptibility or enhanced resistance, depending on the genetic context. For instance, while mutants such as *NahG*, *jai1*, *spr2*, and *def1*-which exhibit impaired salicylic acid (SA) or jasmonic acid (JA) signaling-tend to be more susceptible to various pathogens or herbivores (López-Gresa et al., 2016; Li et al., 2004; Howe et al., 1996), other genetic perturbations-such as *sldmr6-1* or RNAi *SlS5H*-enhance disease resistance by modulating hormone homeostasis or activating defense responses (Thomazella et al., 2021; Payá et al., 2022). Notably, in *SlPLC2* KO plants, both SA- and JA-related gene expressions are downregulated during infection, and yet plants exhibit less susceptibility to *P. infestans*. This contrasts with the established pattern where disruption of hormone signaling or synthesis pathways weakens defense, suggests that SlPLC2 promotes disease not simply by modulating hormone output, but by facilitating pathogen compatibility, positioning it as a *bona fide* susceptibility gene whose inactivation enhances resistance.

Interestingly, *SlPLC2* KO plants also accumulated higher levels of SA under mock conditions (detached leaflets incubated under high humidity), suggesting that *SlPLC2* deficiency may alter responsiveness to environmental or mechanical stimuli. While this observation does not reflect basal physiology, since SA levels in planta are the same as in WT, it provides indirect evidence that *SlPLC2* influences the sensitivity or threshold of hormonal signaling pathways even in the absence of infection. Notably, levels of other hormones such as abscisic acid (ABA) and indole-3-acetic acid (IAA) remained unchanged in the KO lines, reinforcing the idea that SlPLC2 selectively affects defense-related hormonal circuits.

Altogether, these findings indicate that *SlPLC2* KO plants exhibit an overall attenuation of SA and JA signaling during *P. infestans* infection. This hormonal phenotype suggests that SlPLC2 plays a positive regulatory role in the activation of these pathways. The impaired transcriptional induction of SA- and JA-related genes in the absence of *SlPLC2* underscores its functional contribution to the immune hormone signaling network.

### ROS and immune defense

*SlPLC2* KO lines exhibited impaired defense responses upon *P. infestans* infection, including reduced accumulation of ROS and diminished callose deposition suggesting that SlPLC2 contributes directly to the proper activation of immune responses. These responses are essential for initial immune activation and cell wall reinforcement (Mittler, 2017; Kadota et al., 2014). AtPLC2 interacts with RBOHD, regulating ROS production (D’Ambrosio et al., 2017). Possibly SlPLC2 operates within a conserved lipid-mediated signaling cascade. Although only ROS were directly measured in this study and other components were not assessed, the signaling module appears conserved, suggesting that SlPLC2 may coordinate immune activation, including Ca²⁺ and MAPK cascades, and gene expression. This interpretation is further supported by the reduced callose deposition observed in *SlPLC2* KO plants, a defense response that functions downstream of ROS accumulation, indicating that immune signaling is attenuated. In line with this, the expression of defense-related genes associated with SA and JA pathways are also diminished in the knockout lines.

Interestingly, despite reduced ROS accumulation, *SlPLC2* KO plants were less susceptible to *P. infestans*. A similar trend was reported in transgenic potato plants overproducing L-ascorbate, which exhibited reduced hydrogen peroxide levels yet showed delayed disease progression and milder necrotic symptoms upon *P. infestans* infection (Chung et al., 2019). This apparent contradiction highlights the complex role of ROS in interactions with hemibiotrophic pathogens. Rather than acting as purely protective agents, ROS must be tightly regulated, as their over- or underproduction can compromise defense (Torres et al., 2005; Camejo et al., 2016; Qi et al., 2017; Smirnoff & Arnaud, 2019). In this context, SlPLC2 may modulate the intensity and spatial distribution of oxidative bursts during infection. Other mechanisms, such as tomato long non-coding RNAs (lncRNAs) that fine-tune ROS output by regulating redox-related gene expression during *Phytophthora* infection, illustrate additional layers of redox control (Cui et al., 2017). Disruption of *SlMYBS2*, a gene previously associated with defense responses, results in increased susceptibility to *P. infestans*, characterized by larger lesion areas, extensive cell death, and enhanced accumulation of ROS (Liu et al., 2021). These distinct but complementary pathways highlight the multifaceted regulation of ROS homeostasis during host-pathogen interactions. Supporting this, *NbPLC2*-silenced *N. benthamiana* plants exhibited impaired ROS and callose responses to bacterial elicitors (Kiba et al., 2020). However, in that case, the suppression of *NbPLC2* led to increased susceptibility to *Pseudomonas syringae*, contrasting with the enhanced resistance observed in *SlPLC2* KO lines. This discrepancy may reflect differences in the pathogens used (bacteria vs. oomycetes), or functional divergence among PLC paralogs. Together, these observations support a role for SlPLC2 in modulating immune responses during the onset of *P. infestans* infection. In particular, SlPLC2 appears to influence the oxidative burst and cell wall-associated defenses that shape the outcome of early colonization.

### SlPLC2 localizes in host derived membrane surrounding infection vesicles and contributes to lipid remodeling

We observed a significantly lower number of infection vesicles, each delimited by a membrane interface analogous to the EHM, in SlPLC2 KO plants at 1 dpi. These vesicles represent the first physical interfaces between *P. infestans* and host cells, and their reduced formation may stem directly from the absence of SlPLC2 activity. Fewer infection vesicles could attenuate immune outputs in KO plants, resulting in a lower degree of pathogen proliferation together with a reduced host immune response. Thus, we studied the SlPLC2 localization during infection. Under non-infected conditions, SlPLC2-GFP localizes to the plasma membrane. However, during *P. infestans* infection, it is also localized at the membrane surrounding the infection vesicles. These vesicles are pathogen-induced host compartments that function as specialized interfaces for effector delivery and immune suppression (Bozkurt et al., 2011). Infection vesicles represent one of the earliest host-pathogen interfaces established after appressorium penetration, preceding haustorium formation and serving as an initial site of effector delivery and hyphal emergence. Despite their importance in early colonization, host-derived proteins have rarely been reported in these structures. Our observation of SlPLC2-GFP in infection vesicles thus constitutes the first report of a host-encoded enzyme localized to this compartment, highlighting a previously unexplored layer of host membrane remodeling at the onset of *P. infestans* invasion. Given the current constraints of our imaging system, we could not resolve SlPLC2 localization at the haustoria. Further studies are planned to address this limitation. Nevertheless, the presence of SlPLC2 in infection vesicles therefore opens a new perspective on host phosphoinositide signaling and protein localization during the earliest stages of pathogen invasion, before haustoria differentiation is established.

Recent studies have revealed that the EHM, exhibits a phosphoinositide and protein composition that diverges markedly from that of the plasma membrane, likely due to selective vesicular trafficking and membrane remodeling events during pathogen colonization (Bozkurt & Kamoun, 2020). Rausche et al. (2021) characterized StIPP, a PI(4,5)P₂ 5-phosphatase that hydrolyzes PI(4,5)P₂ to PI4P and localizes to the EHM during *P. infestans* infection. While this enzyme is transcriptionally induced by PAMPs and localizes to infection sites, silencing StIPP in potato did not lead to increased susceptibility or resistance, suggesting that its contribution to defense is subtle or redundant under native conditions. In Arabidopsis, Qin et al. (2020) showed that PI(4,5)P₂ accumulates at the EHM during infection by *Erysiphe cichoracearum*, and that pip5k1 pip5k2 double mutants-lacking two PI4P 5-kinases responsible for PI(4,5)P₂ synthesis-display strong resistance to *E. cichoracearum* and *Albugo candida*. In addition, Shimada et al. (2019) used an estradiol-inducible system to transiently express PIP5K3-GFP and demonstrated that increased PI(4,5)P₂ accumulation at the extra-invasive hyphal membrane (EIHM) correlates with enhanced susceptibility to *Colletotrichum*. In contrast, PI(4,5)P₂ enrichment was not observed at the EHM during infection by *Hyloperonospora Arabidopsis* or *Golovinomyces orontii*, indicating that this lipid remodeling is specific to the interaction. In *N. benthamiana* roots colonized by the mutualistic fungus *Funneliformis mosseae,* Guyon et al. (2025) showed that PI4P, rather than PI(4,5)P₂, accumulates at the EHM during *Phytophthora palmivora* infection correlating with enhanced resistance. Notably, the resistant state and altered phosphoinositide landscape were dependent on prior mycorrhizal colonization, indicating that mutualists can reprogram host membrane identity to restrict pathogen invasion. In *Lotus japonicus* during symbiotic infection by *Mesorhizobium loti* (Akamatsu et al., 2025) mutants impaired in phosphoinositide metabolism (pip5k4, pip5k6, plp4) exhibited reduced PI(4,5)P₂ content and enhanced infection thread formation. These findings suggest that, unlike in pathogenic interactions with filamentous microbes, PI(4,5)P₂ acts as a negative regulator of infection in *L. japonicus*, serving as a molecular brake to prevent over-colonization. That evidence is consistent with findings by Guyon et al. (2025), who showed that PI4P accumulation at the EHM correlates with enhanced resistance in mycorrhiza-primed roots during *P. palmivora* infection, and with observations by Ivanov & Harrison (2018), who reported the accumulation of PI4P and PI(4,5)P₂ in distinct domains of the periarbuscular membrane during arbuscular mycorrhizal symbiosis. This highlights the context-dependent role of phosphoinositides, which can either promote or restrict microbial accommodation depending on whether the interaction is pathogenic or symbiotic. Although PA distribution was not addressed in the mycorrhiza study, one could speculate that the absence or scarcity of PA, that is produced downstream DAG formation by SlPLCs, might prevent phosphoinositide from ‘diverting’ towards pathogen-favorable membrane remodeling, thereby maintaining a symbiotic-compatible interface.

SlPLC2 localizes to infection vesicles, where we detected the presence of PI4P and PI(4,5)P₂. This spatial coincidence suggests that SlPLC2 may act on infection vesicle membranes containing phosphoinositide during *P. infestans* colonization, modulating their composition and function. Notably, the relative distribution of these lipids in infection vesicles differs from that observed at the plasma membrane, further supporting the notion that these vesicles represent a distinct and pathogen-induced membrane domain. Our findings position SlPLC2 as a modulator of membrane lipid composition at infection vesicles during *P. infestans* colonization. By hydrolyzing PI4P and PI(4,5)P₂, SlPLC2 promotes the depletion of these lipids from membranes, triggering the generation of DAG and IP₂/IP₃ and possibly initiating downstream events that support vesicle expansion and structural remodeling. This enzymatic activity likely facilitates the formation and stabilization of membrane compartments required for effector secretion and pathogen progression in the infection vesicle. In barley, silencing of *PLC1* enhances susceptibility to the biotrophic fungus *Blumeria hordei* (Weiß et al., 2025). In the absence of SlPLC2, higher local levels of PI4P and PI(4,5)P₂ may persist at infection sites and its excessive accumulation can disrupt the specialized membrane environments required for infection progression. In line with this, *SlPLC2* knockout plants are less susceptible to *P. infestans*, whereas overexpression of SlPLC2 in *N. benthamiana* increases susceptibility. These results support a model in which SlPLC2 also promotes infection by catalyzing the turnover of phosphoinositide at early host-pathogen interfaces, enabling vesicle biogenesis and effector accommodation.

### Proposed model and concluding remarks

Based on our results and previous studies, we propose a model describing the role of SlPLC2 during *P. infestans* infection (Figure 7). SlPLC2 localizes to the plasma membrane and to the membranes surrounding infection vesicles. At these pathogen-induced membrane domains, SlPLC2 may hydrolyze PI4P and PI(4,5)P₂, generating the second messengers IP₂/IP₃ and PA (derived from the DAG phosphorylation). This localized lipid turnover could facilitate membrane remodeling and the accommodation of pathogen structures, promoting colonization. In parallel, at the plasma membrane, SlPLC2 may participate in immune signaling by generating IP₂/IP₃/PA upon pathogen recognition. These messengers trigger calcium influx and ROS production via RBOHD, and downstream regulation of SA- and JA-defense responses, linking SlPLC2 to both membrane remodeling and immune signal transduction. Although the two processes are represented separately in our model, we cannot exclude the possibility that the signaling events illustrated at the plasma membrane also take place at the host-derived membrane enclosing the infection vesicle. It would be interesting to see, for instance RBOHD localization at the host-derived membrane.

**Figure 7.**
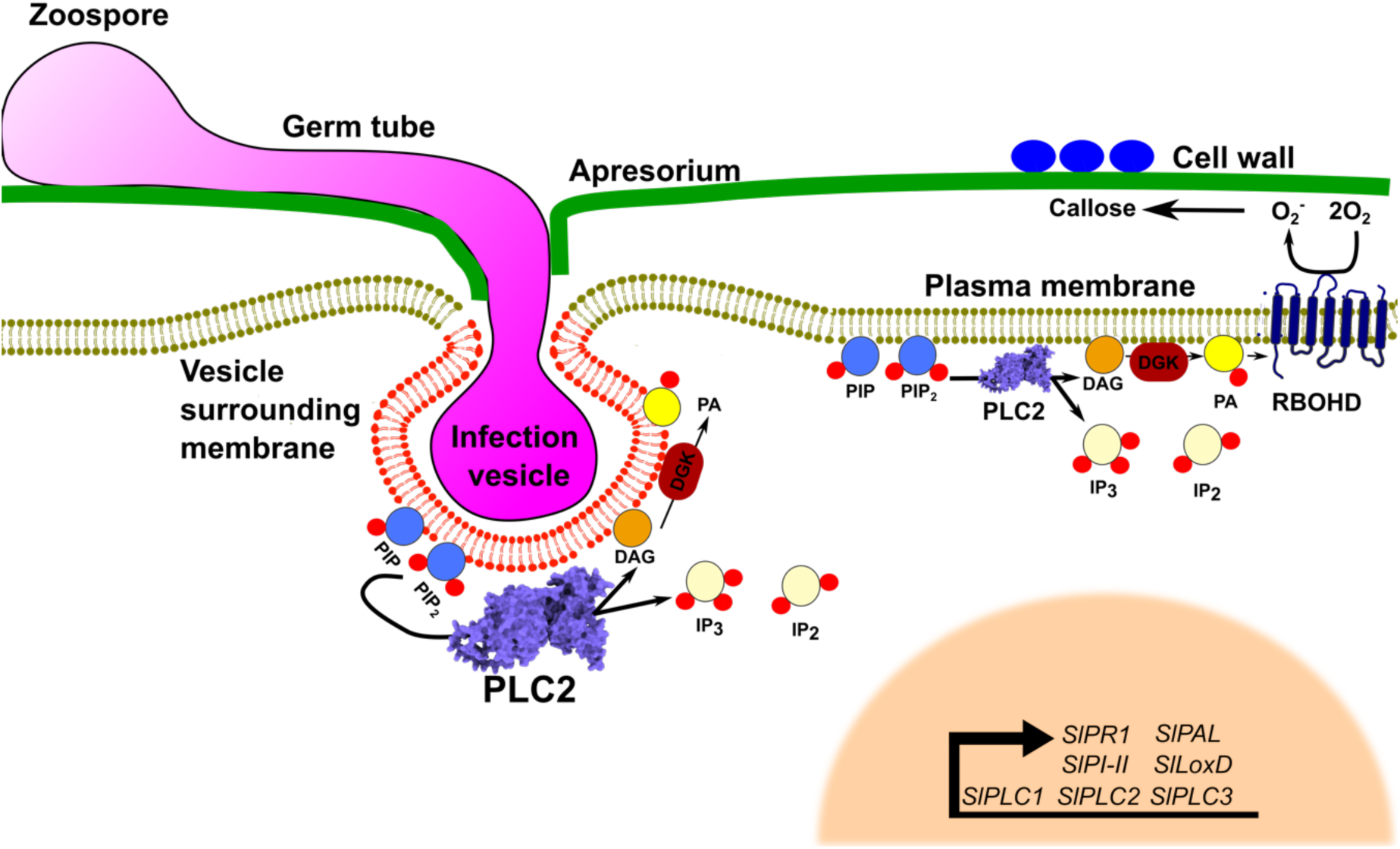
Proposed model of *P. infestans* infection and the role of SlPLC2. Upon contact with the host surface, *P. infestans* establishes infection vesicles as the initial intracellular interfaces. These pathogen-induced compartments are associated with early immune responses such as reactive oxygen species (ROS) production and callose deposition. SlPLC2 localizes both to the plasma membrane and to membranes surrounding the vesicle membrane, where it hydrolyzes phosphoinositides PI4P and PI(4,5)P₂, generating second messengers that may modulate immune signaling and vesicle remodeling. In the image, the membrane surrounding the infection vesicle is shown in red and the host plasma membrane is shown in green.

This work represents one of the first reports of a host lipid-signaling enzyme accumulating at *P. infestans*-induced infection vesicles. *SlPLC2* exemplifies a class of dynamically localized susceptibility factors whose lipid signaling activity is intimately linked to pathogen accommodation. Unraveling how such host components contribute to vesicle formation and immune modulation will expand our understanding of host-pathogen interactions and open new avenues for engineering durable resistance in solanaceous crops.

## Materials and Methods

### Plant and oomycete material

Tomato plants (*Solanum lycopersicum* cv. Moneymaker, MM-Cf0) and CRISPR/Cas9-generated knockout lines KO-6 and KO-17 (Perk et al., 2023) were grown from seeds in commercial Growmix substrate under controlled environmental conditions (16 h light / 8 h dark photoperiod, 25 °C, 70% relative humidity). Plants were used at 6 weeks of age. For each genotype, five plants were analyzed per biological replicate, and experiments were conducted in triplicate unless otherwise stated.

*Nicotiana benthamiana* plants were also grown from seed in Growmix under the same environmental conditions and used at 4 weeks of age for transient expression assays.

The *P. infestans* isolate used in this study, designated PinSL-19, was previously obtained by (Juarez et al 2023). PinSL-19 was identified as belonging to the EU_2_A1 genotype, the dominant clonal lineage currently circulating in Argentina, Chile, and Brazil (Juarez et al 2023).)

### *P. infestans i*nfection assay

*P. infestans* isolate PinSL-19 was maintained on Rye Sucrose Agar (RSA) as described by Caten & Jinks (1968), and zoospores were isolated following Tooley et al. (1986). Six-week-old tomato leaves were detached and placed in plastic humidity chambers (sealed trays covered with plastic film). For each genotype, ten leaflets from five individual plants (two leaves per plant) were used per biological replicate, and the experiment was performed in triplicate. Each lateral leaflet (excluding the central leaflet) was inoculated on the abaxial with two droplets of 10 µL of 2 × 10^5^ zoospore/mL suspension using a multichannel repeating pipette. Control leaves were inoculated in parallel with sterile water. Inoculated leaves were incubated at 25 °C with high humidity, and infection progression was evaluated at 6 days post-inoculation (dpi) by measuring lesion area per inoculation point using ImageJ software.

### RNA extraction and RT-qPCR gene expression analysis

Total RNA was extracted from tomato leaves collected at 0, 1, 2, 4, and 6 days post-inoculation (dpi) with *P. infestans*, using the TRIzol reagent according to the manufacturer’s instructions (Invitrogen). RNA integrity was assessed by agarose gel electrophoresis, and concentration and purity were evaluated using a Nanodrop spectrophotometer (Thermo Fisher Scientific), accepting samples with A260/A280 and A260/A230 ratios above 1.80.

One microgram of total RNA was reverse-transcribed using M-MLV reverse transcriptase (Invitrogen) and oligo(dT) primers. Quantitative PCR was performed using a 1:5 dilution of the cDNA as template and the Fast Universal SYBR Green Master Mix (Roche, Mannheim, Germany), on a Step One Plus Real-Time PCR System (Applied Biosystems). Primer sequences for *SlPLC1-6*, *SlPR1a, SlPI-II, SlACTIN,* and *PiEF1-α* were previously reported (Gonorazky et al., 2016) and *PiEF1-α*, listed in Table S1. Relative transcript levels were calculated using LinRegPCR software (Ramakers et al., 2003), which estimates individual amplification efficiencies. Expression values were normalized against the *SlACTIN* reference gene. Each sample was analyzed in two technical replicates, and biological replicates ranged from 4 to 12, as specified in the corresponding figure legends.

### Sporulation Assay in Tomato Leaf Discs

Tomato wild-type (WT) plants and SlPLC2 knockout lines KO-6 and KO-17 were grown as described above. Leaf discs (10 mm in diameter) were excised from the third and fourth fully expanded leaves and placed individually in 24-well plates containing 1 mL of water agar (0.8%). Each disc was inoculated with a 10 µL droplet of a *P. infestans* 2 × 10^4^ zoospore/mL suspension (strain PinSL-19). Plates were incubated in a growth chamber at 20°C with a 16 h light / 8 h dark photoperiod. At 2,3,4,5 and 6 dpi, leaf discs were transferred to 1.5 mL microcentrifuge tubes containing 0.5 mL of phosphate-buffered saline (PBS) supplemented with 1% formaldehyde. Tubes were vortexed thoroughly, and sporangia concentration was quantified using a Neubauer counting chamber. Data were expressed as the number of sporangia per milliliter.

### Phytohormone quantification

The levels of salicylic acid (SA), jasmonic acid (JA), and methyl jasmonate (MeJA) were quantified in tomato leaves at 6 days post-inoculation (dpi) with *P. infestans*. For each sample, the four lateral leaflets from a single infected leaf were pooled, flash-frozen in liquid nitrogen, and ground to a fine powder. Five biological replicates were processed per treatment.

Hormone extraction was performed following Otaiza-González et al. (2022), with modifications. Approximately 50-100 mg of powdered tissue were homogenized in 500 µL of a 2:1:0.002 (v/v/v) solution of 1-propanol:water:concentrated HCl, shaken for 30 min at 4 °C, and mixed with 1 mL of dichloromethane (CH₂Cl₂). Samples were vortexed, incubated for another 30 min at 4 °C, and centrifuged at 13,000 × g for 5 min. The lower organic phase (∼1 mL) was transferred to a new vial and evaporated under a nitrogen stream. Dried residues were resuspended in 250 µL of 1:1 (v/v) methanol:water (both HPLC grade) containing 0.1% formic acid.

Phytohormones were analyzed by LC-MS/MS using a Waters Xevo TQ-S micro mass spectrometer equipped with an Acquity UPLC H-Class system, autosampler, and BEH C18 column (1.7 µm, 2.1 × 50 mm, Waters). The mobile phase consisted of solvent A (water + 0.1% formic acid) and solvent B (methanol + 0.1% formic acid), with a flow rate of 0.25 mL/min. The gradient started at 40% B, increased linearly to 100% over 3 min, and was maintained for 0.5 min. Mass detection was performed in positive electrospray ionization (ESI+) mode using MassLynx software.

Quantification was carried out by external calibration with commercial standards (Sigma-Aldrich) for SA, JA, and MeJA. Results were expressed as nanograms of phytohormone per milligram of fresh weight.

### Detection of H₂O₂ in *P. infestans*-infected leaves

Hydrogen peroxide (H₂O₂) accumulation in response to *P. infestan*s was detected using 3,3-diaminobenzidine (DAB) staining as described by Thordal-Christensen et al. (1997), with minor modifications. Inoculated tomato leaflets were harvested at 1-day post-inoculation (dpi), cleared in 96% ethanol until tissue was translucent, and then incubated in a solution of 15 mg L⁻¹ DAB dissolved in 0.5 M sodium acetate buffer (pH 5.5) for 1 to 5 hours at room temperature. DAB reacts with H₂O₂ in the presence of endogenous peroxidases to form a brown precipitate. After staining, leaflets were rinsed in water and mounted for analysis. Stained areas were imaged and quantified point-by-point using ImageJ software, and signal intensity was expressed as mean gray value per mm². Each treatment included ten leaflets from five plants (as in the infection assay), with three independent biological replicates.

### Callose deposition

Callose accumulation was assessed at 2 dpi in *P. infestans*-inoculated tomato leaflets. From each leaflet, eight 5-mm discs were collected, each corresponding to an individual inoculation site. Samples were cleared in 96% ethanol until completely transparent, then washed in 0.07 M phosphate buffer (pH 9.0) and incubated for 2 h in the same buffer containing 0.01% (w/v) Aniline Blue. Stained discs were mounted in glycerol and observed under a Nikon C1siR confocal laser scanning microscope equipped with a UV filter (excitation 365/10 nm; emission 460/50 nm), using a 10× objective. Callose deposits were visualized as fluorescent spots, and their number was quantified using ImageJ software (Schneider et al., 2012). For each biological replicate (n = 3), twenty measurements were taken per sample, averaging callose dot counts per field from one representative disc per leaflet.

### Quantification of infection vesicles

To quantify infection vesicles, tomato leaf was drop-inoculated on the abaxial surface with a zoospore suspension of Td 88069 (2 × 10⁵ zoospores/mL) as for PinSL-19 infection assay. At 1 day post-inoculation (dpi), 1 cm diameter disks were excised using a sterile cork borer. A total of 24 disks per biological replicate were collected and immediately mounted in water on glass slides. Vesicles were visualized in epidermal cells using an epifluorescence microscope (Zeizz axio imager.A2) Camera (AxioCam HRc). At least three fields per disk were imaged to quantify the number of vesicle-containing cells. Three biological replicates were analyzed.

### Gene expression analysis of *PiAvrblb2* and *PiEF1-α*

For expression analyses, tomato leaflets were infected with *P. infestans* and sampled at the indicated time points. From each leaflet, a 1 cm diameter disk was excised using a cork borer and immediately frozen in liquid nitrogen. For each biological replicate (n = 4), total RNA was extracted from a pool of 8 disks using the same extraction protocol. Transcript levels of the biotrophic effector gene *PiAvrblb2* and the constitutive P. infestans gene *PiEF1-α* were measured by RT-qPCR using gene-specific primers (Supplemental Table S1). Relative expression of *PiAvrblb2* was calculated by first normalizing to the *P. infestans* gene *PiEF1-α* and then relativizing to the expression levels detected in plate-grown *P. infestans*. *PiEF1-α 1*expression was normalized to plant constitutive gene *SlACTIN*.

### Transient expression in *Nicotiana benthamiana*

Transient expression assays were performed in 4-week-old *N. benthamiana* plants by *Agrobacterium tumefaciens*-mediated infiltration using the GV3101 strain carrying the binary vector pEAQ (Lomonossoff et al., 2009) with SlPLC2-GFP or GFP alone as a control. Additionally, fluorescently tagged lipid sensors specific for PI4P (mCitrine-2xPH_FAPP_) and PI4P₂ (mCitrine-2XPH_PLC_ were used for colocalization experiments (Simon et al 2014). A. tumefaciens cultures were grown overnight at 28 °C with shaking (200 rpm) in LB medium with appropriate antibiotics. The cultures were centrifuged at 4000 rpm for 8 minutes and resuspended in freshly prepared MMA infiltration buffer (50 mL: 1 g sucrose [58.4 mM], 0.25 g Murashige and Skoog salts [∼4.3 mM], 0.1 g MES [4.9 mM], and 200 µM acetosyringone) to a final OD₆₀₀ of 0.2-0.3. Bacterial suspensions were incubated for 1-6 h at room temperature to induce virulence genes.

Approximately 2-3 leaves from two individual plants were infiltrated per construct, and five independent biological replicates were performed. In some assays, transient expression was followed by infection of detached leaves with Td88069 under dark conditions (18 °C, 60% relative humidity), as described by Bozkurt et al. (2014). Fluorescence was visualized 48 hours post-infiltration using a Nikon C1siR confocal laser scanning microscope.

### Statistical analysis

Statistical analyses were performed using generalized linear mixed models (GLMMs), implemented with the lme (from the *nlme* package) or glmer (from the *lme4* package) functions in R (version 3.1; R Foundation for Statistical Computing). Genotype (WT or KO lines) was included as a fixed effect in all models, while biological replicate and, when applicable, humidity chamber was treated as random effects.

For time-course experiments (e.g., gene expression and hormone quantification), both genotype and time point were modeled as fixed effects, and their interaction was tested. In these cases, post hoc comparisons were used to identify significant differences across all genotype × time combinations, and results are shown using letters to indicate statistically distinct groups.

In other experiments where only genotype was considered as a fixed effect (e.g., ROS production, callose deposition, infection vesicle quantification), statistically significant differences relative to the WT control are indicated with asterisks, based on Dunnett’s post hoc tests.

For the sporulation assay, statistical comparisons were restricted to within each time point, since a time-dependent increase in spore number is biologically expected and not informative for genotype effects. Data did not meet the assumptions of normality and homoscedasticity, so a Kruskal-Wallis test was used followed by pairwise Wilcoxon tests. Statistically significant differences relative to WT are indicated with asterisks (* = p ≤ 0.05; ** = p ≤ 0.01; *** = p ≤ 0.001) based on the Wilcoxon test.

Model assumptions for GLMMs were assessed by visual inspection of residuals (Q-Q plots and residual vs. fitted plots). Depending on the residual distribution and dispersion, either Gaussian or Gamma error distributions were applied.

### Confocal microscopy

All confocal images were acquired using a Nikon C1siR laser scanning confocal microscope equipped with a UV excitation filter (Ex 365/10 nm, Em 460/50 nm) and Plan Fluor objectives (40×). Imaging was performed using Nikon EZ-C1 software. Laser intensity, detector gain, and offset were kept constant across all samples within each experiment. For Aniline Blue staining, excitation was performed with a 405 nm laser line, and emission was collected between 450-480 nm. For GFP-based constructs, excitation was set at 488 nm and emission collected between 500-530 nm. All images were acquired under identical acquisition settings per experiment.

### Western blot analysis

Proteins were extracted by homogenizing 4-5-week-old leaf tissue in protein extraction buffer [100 mM NaPi pH 7.5, 150 mM NaCl, 1 mM EDTA, and protease inhibitor cocktail (SIGMA)] at a 1:1 volume ratio. Samples were centrifuged for 10 min at 10,000 g, and total protein concentration was determined in the supernatant. Proteins were purified via affinity chromatography using a His-tag purification column or directly prepared by mixing leaf powder with SDS-PAGE sample buffer without prior extraction. Protein samples were separated on 10% SDS-polyacrylamide gels and transferred to nitrocellulose membranes. Membranes were stained with Ponceau S to verify equal loading and subsequently incubated overnight in PBS-T containing anti-GFP polyclonal antibody (1:2000). After three washes with PBS-T, membranes were incubated with alkaline phosphatase-conjugated anti-rabbit IgG secondary antibody and developed following the manufacturer’s instructions (SIGMA).

### Agarose gel electrophoresis of PCR products

PCR products were obtained using gene-specific primers and cDNA synthesized from total RNA extracted from infected leaf tissue. Amplification reactions were loaded onto 1.5% agarose gels DNA bands were visualized under UV illumination and imaged using a gel documentation system. A 100 bp DNA ladder was used as a size marker.

## Acknowledgments

We thank our colleagues at the Instituto de Investigaciones Biológicas (IIB-CONICET-UNMdP) for discussions and technical support. We also acknowledge the use of ChatGPT (OpenAI, GPT-5) to improve English (writing and grammar) in the preparation of this manuscript. All scientific content, data interpretation, and conclusions are the sole responsibility of the authors.

## Funding

This work was supported by the National Scientific and Technical Research Council of Argentina (CONICET), Agencia Nacional de Promoción Científica y Tecnológica (ANPCyT, PICT grants), and Universidad Nacional de Mar del Plata (UNMdP).

## Conflict of Interest

The authors declare no conflict of interest.

## Data Availability

All data supporting the findings of this study are available within the manuscript and its supplementary materials. Additional datasets generated during this study, including raw qPCR data, hormone measurements, and confocal images, are available from the corresponding author upon reasonable request.

## Plant Materials

Tomato (*Solanum lycopersicum* cv. Moneymaker, MM-Cf0) wild-type plants and CRISPR/Cas9-generated *SlPLC2* knockout lines (KO-6 and KO-17) were used in this study. Knockout lines were previously described in Perk et al. (2023). Seeds are available from the authors upon request, subject to institutional and national regulations regarding the distribution of genetically modified organisms.

## Author contributions

E.A.P. conceived and designed experiments, performed all assays except sporulation, conducted statistical analyses, interpreted results, and wrote the manuscript. J.M.D. performed confocal and epifluorescence microscopy and assisted in data analysis and interpretation. I.C. contributed to the construction and generation of transgenic lines and assisted in infection assays. L.R. performed calose deposition assays, analyzed the results, and carried out immunoblotting. M.E.S. and M.J. conducted sporulation assays, provided *P. infestans* isolates, contributed expertise for microscopy experiments, and assisted in literature review and interpretation of results. P.V., M.T., and V.M. provided equipment and support for hormone quantification. A.M.L. conceived and supervised the project, contributed to experimental design and discussion of results, and is the corresponding author. All authors reviewed and approved the final version of the manuscript.

## Supplemental Figures

**Supplementary Figure S1.**
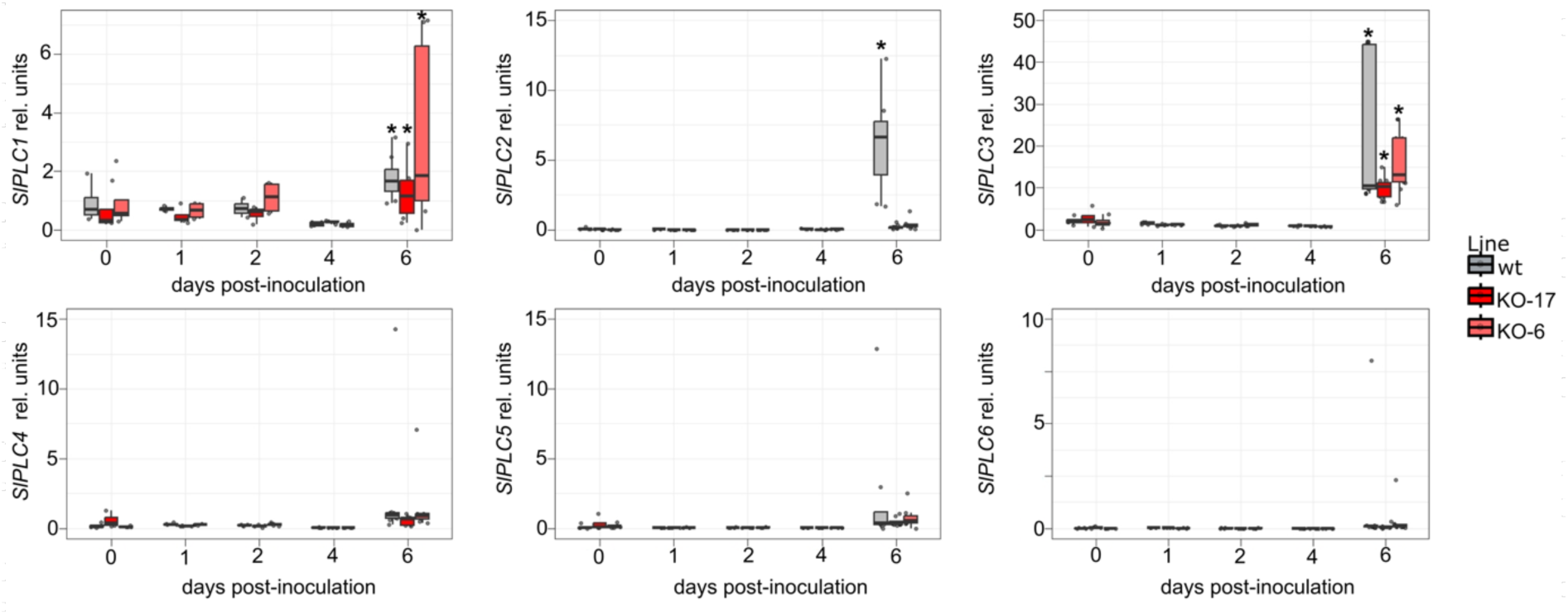
Expression of *SlPLC* gene family members during *P. infestans* infection. Time-course RT-qPCR analysis of *SlPLC1-6* transcript levels in tomato leaflets inoculated with *P. infestans* (10 µl droplet of a 2×10^5^ zoospore per mL). Samples were collected at 6 days post-inoculation (dpi) or from non-infected controls. Transcript levels were normalized to *SlACTIN* and expressed relative to non-infected samples. Data represent mean ± SE of four biological replicates (mock) and eight (infected). Asterisks indicate statistically significant differences compared to mock (Dunnett’s test, *P* < 0.05).

**Supplementary Figure S2.**
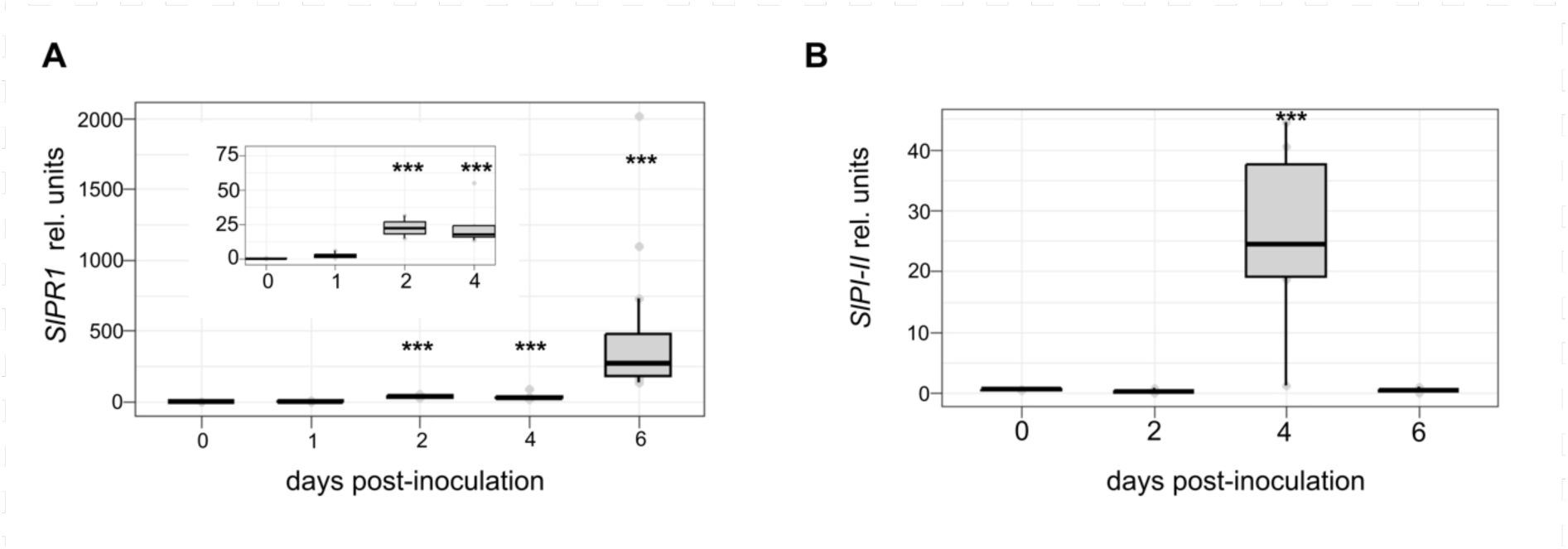
Salicylic acid and Jasmonic acid pathway marker expression during *P. infestans* infection. **(A)** Time-course RT-qPCR analysis of *SlPR1a* transcript levels in wild-type (WT) tomato plants at 2, 4, and 6 dpi. **(B) (A)** Time-course expression of *SlPI-II* in WT plants at 2, 4, and 6 dpi, measured by RT-qPCR. Transcript levels were normalized to *SlACTIN*. Data represent mean ± SE of 4-12 biological replicates per time point.

**Supplementary Figure S3.**
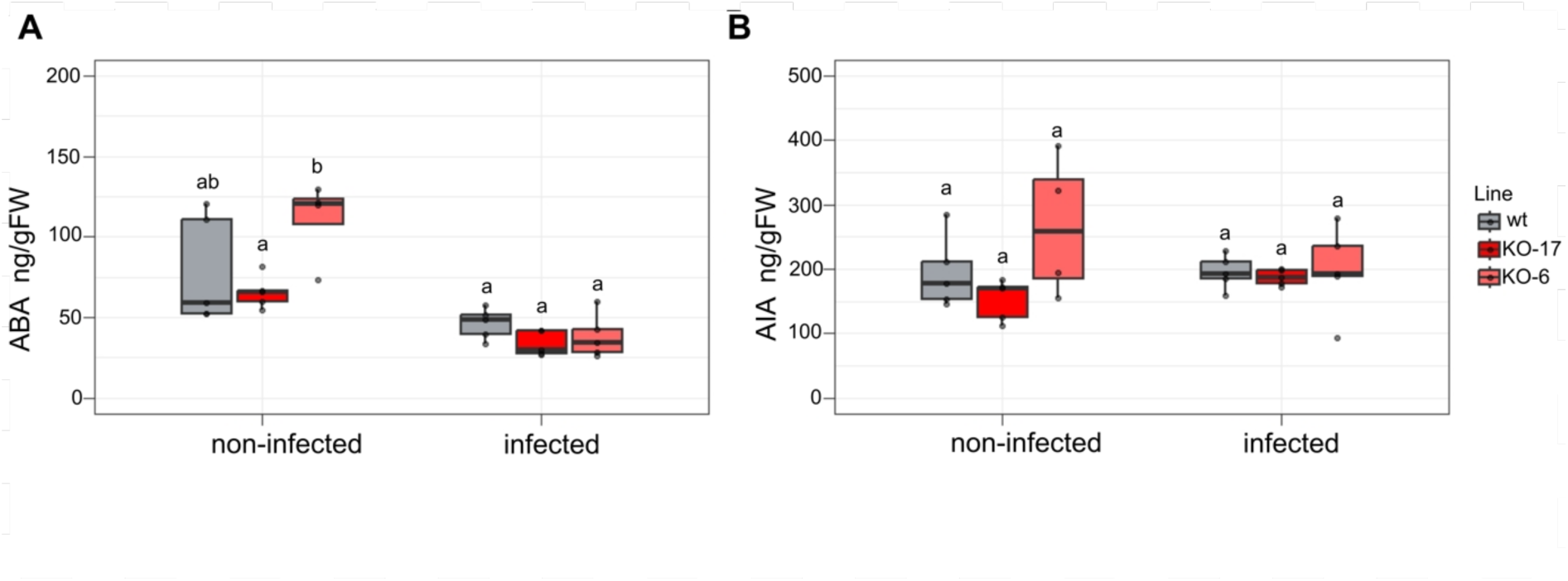
ABA and IAA remain unchanged in SlPLC2 knockout lines during *Phytophthora infestans* infection. WT and SlPLC2 knockout tomato lines (KO-6 and KO-17) were inoculated with *P. infestans* zoospores by applying two 10 µL droplets of a 2 × 10⁵ zoospores/mL suspension on either side of the central vein of each leaflet. **(A-B)** Quantification of endogenous hormone levels by LC-MS/MS in aerial tissues at 6 dpi in infected or non-infected plants. Hormone concentrations were expressed as nanograms per milligram of fresh weight. **(A)** Abscisic acid (ABA), **(B)** Indole-3-acetic acid (IAA). Data represent the mean ± SE of biological replicates (n = 5). No statistically significant differences were detected between genotypes under either condition (Dunnett’s test, P > 0.05).

**Supplementary Figure S4.**
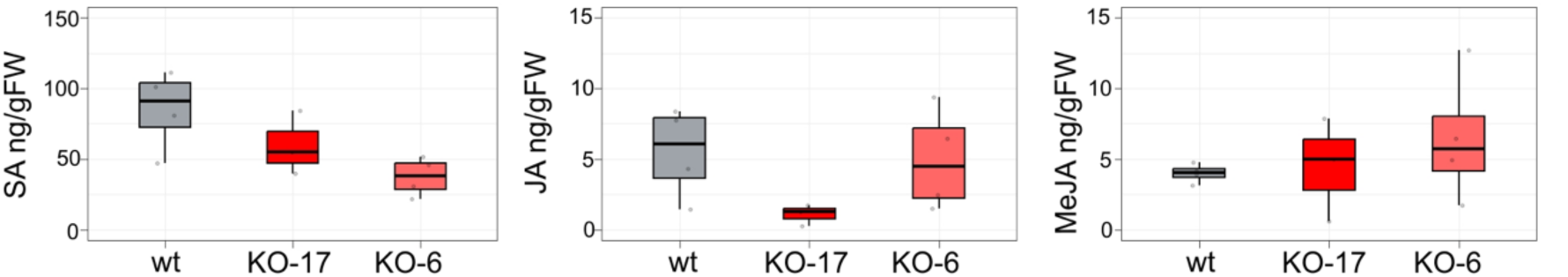
Basal levels of SA, JA, and MeJA are not altered in SlPLC2 knockout lines under non-infected conditions. Detached leaflets from WT and SlPLC2 knockout tomato lines (KO-6 and KO-17) were processed immediately after excision, without any pathogen inoculation. **(A-C)** Quantification of endogenous hormone levels by LC-MS/MS in aerial tissues. Hormone concentrations were expressed as nanograms per milligram of fresh weight. **(A)** Salicylic acid (SA), **(B)** Jasmonic acid (JA), **(C)** Methyl jasmonate (MeJA). Data represent the mean ± SE of biological replicates (n = 5). No statistically significant differences were observed between genotypes (Dunnett’s test, P > 0.05).

**Supplementary Figure S5.**
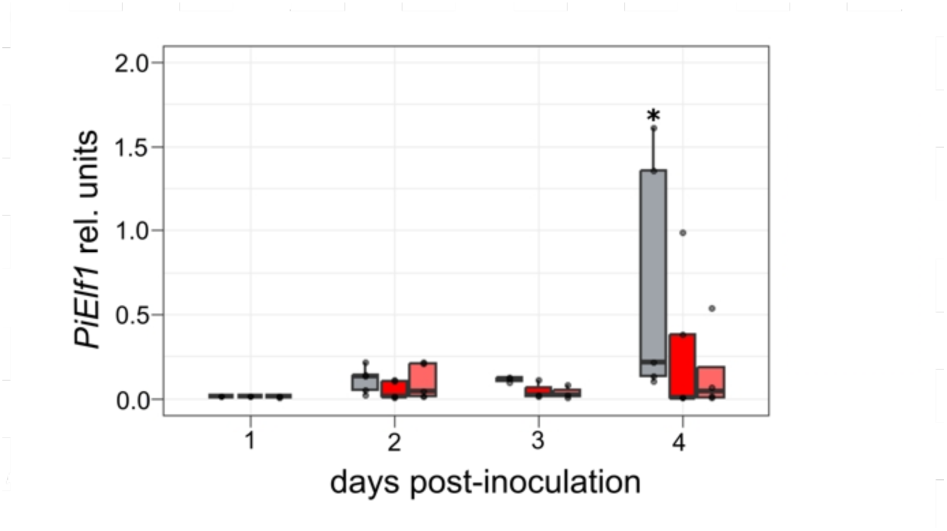
Quantification of *Phytophthora infestans* biomass in tomato leaflets over time. Relative expression levels of the *P. infestans* elongation factor gene (*PiEF1-α*) were measured by RT-qPCR at 1, 2, 3, and 4 dpi in WT and KO tomato lines. Data were normalized to *SlACTIN* and expressed in arbitrary units. Box plots show median, interquartile range, and outliers (n ≥ 4). Asterisks indicate statistically significant differences compared to WT at each time point (ANOVA followed by Tukey’s post-hoc test, *P* < 0.05).

**Supplementary Figure S6.**
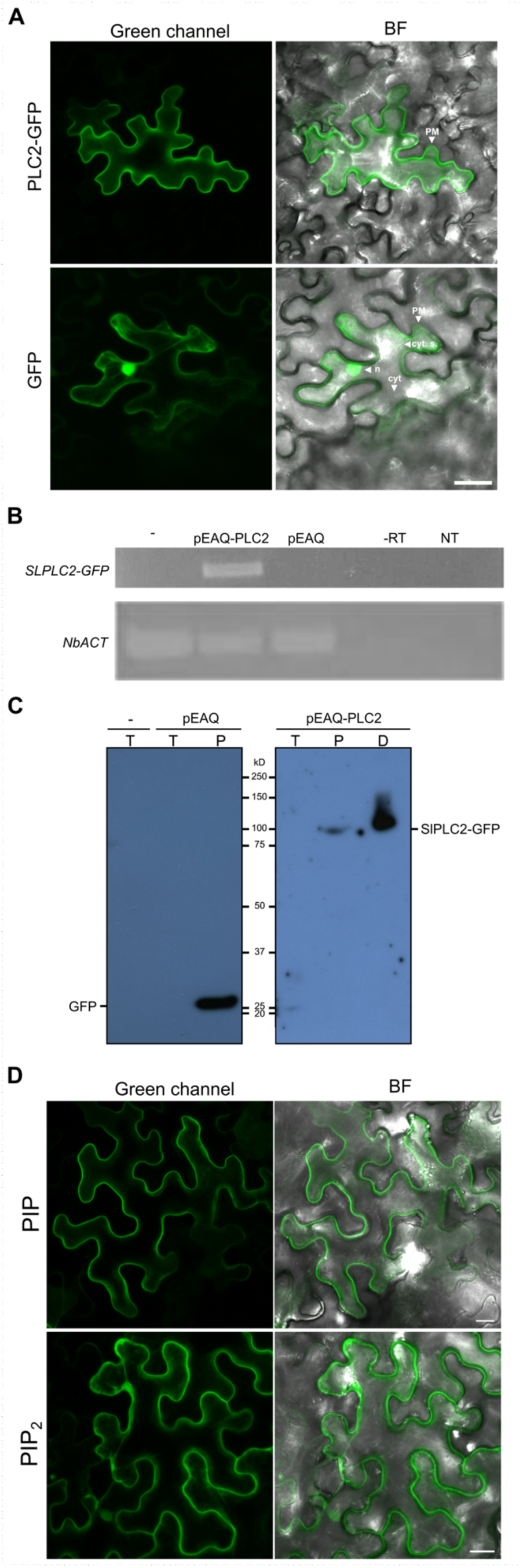
Localization and validation of SlPLC2-GFP expression in *Nicotiana benthamiana*, and distribution of lipid biosensors under basal conditions. All experiments were conducted under non-infected (basal) conditions. **(A)** Confocal microscopy images showing subcellular localization of SlPLC2-GFP in epidermal cells. SlPLC2-GFP localizes predominantly to the plasma membrane. In contrast, free GFP displays diffuse cytosolic and nuclear localization. PM: plasma membrane; n: nucleus; cyt. s.: cytosolic strands; cyt: cytosol. Scale bar = 30 µm. **(B)** RT-PCR detection of the *SlPLC2*-GFP fusion transcript at 2 days post-infiltration confirms transcriptional expression. **(C)** Western blot analysis using anti-GFP antibody confirms protein-level expression of SlPLC2-GFP. Proteins were extracted from 4-5-week-old leaf tissue, separated on 10% SDS-PAGE, and transferred to nitrocellulose membranes. Ponceau S staining was used to verify equal loading. Membranes were incubated with alkaline phosphatase-conjugated anti-rabbit IgG secondary antibody. The positions of molecular weight markers (kDa) are indicated on the left. T: Total, P: Purified with His-tag column, D: Direct. **(D)** Confocal microscopy images showing the localization of PI4P and PI(4,5)P₂ biosensors in *N. benthamiana* epidermal cells. The PI4P biosensor (mCitrine-2xPH(FAPP1)) exhibits strong plasma membrane localization, while the PI(4,5)P₂ biosensor (mCitrine-2xPH(PLC)) displays both plasma membrane and cytosolic distribution. Images were acquired at 2 days post-infiltration. Scale bar = 20 µm.

**Supplementary Figure S7.**
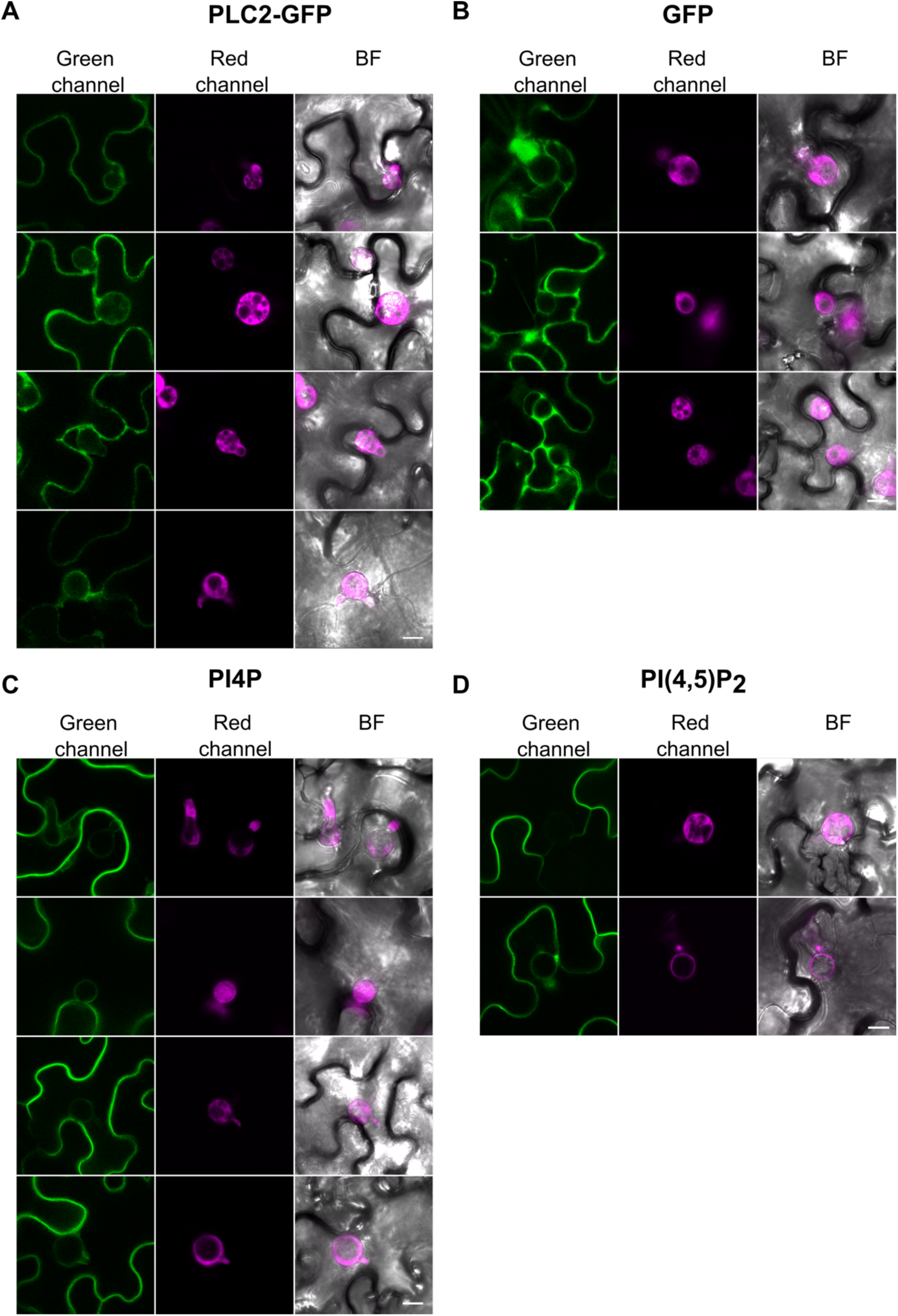
Subcellular localization of SlPLC2-GFP, free GFP, and phosphoinositide biosensors during *Phytophthora infestans* infection. Confocal microscopy images of *Nicotiana benthamiana* epidermal cells at 2 days post-infiltration and 1-day post-inoculation with *P. infestans*. Leaf sectors were co-infiltrated with *Agrobacterium tumefaciens* carrying the indicated constructs and later inoculated with *P. infestans* zoospore suspension (2 × 10⁵ zoospores/mL; 10 µL per site). Whole-cell projections are shown to highlight subcellular distribution and vesicular localization. **(A)** SlPLC2-GFP: Plasma membrane signal with additional accumulation in infection-associated vesicular structures. **(B)** Free GFP: Diffuse cytosolic and nuclear signal, no specific enrichment in vesicles. **(C)** PI4P biosensor (mCitrine-2xPH(FAPP1)): Localizes to the plasma membrane and shows focal accumulations at infection sites. **(D)** PI(4,5)P₂ biosensor (mCitrine-2xPH(PLC)): Displays plasma membrane and cytosolic localization, with enriched labeling at the membranes surrounding infection vesicles. Scale bar = 20 µm.

## Notes

### Competing Interest Statement

The authors have declared no competing interest.

## Bibliography

Abd-El-Haliem, A. M., & Joosten, M. H. A. J. (2017). Plant phosphatidylinositol-specific phospholipase C at the centre of plant innate immunity. Journal of Integrative Plant Biology, 59, 164–179.

Akamatsu, A., Ishikawa, T., Tanaka, H., Kawano, Y., Hayashi, M., & Takeda, N. (2025). Rhizobial infection-specific accumulation of phosphatidylinositol 4,5-bisphosphate inhibits the excessive infection of rhizobia in Lotus japonicus. bioRxiv.

Avrova, A. O., Boevink, P. C., Young, V., Grenville-Briggs, L. J., van West, P., Birch, P. R. J., & Whisson, S. C. (2008). A novel Phytophthora infestans haustorium-specific membrane protein is required for infection of potato. Cellular Microbiology, 10(12), 2271–2284.

Boevink, P. C., Birch, P. R. J., Turnbull, D., & Whisson, S. C. (2020). Devastating intimacy: The cell biology of plant-Phytophthora interactions. New Phytologist, 228(2), 445–458.

Bozkurt, T. O., & Kamoun, S. (2020). The plant-pathogen haustorial interface at a glance. Journal of Cell Science, 133, jcs237958.

Bozkurt, T. O., Richardson, A., Dagdas, Y. F., Mongrand, S., Kamoun, S., & Raffaele, S. (2014). The plant membrane-associated REMORIN1.3 accumulates in discrete perihaustorial domains and enhances susceptibility to Phytophthora infestans. Plant Physiology, 165(3), 1005–1018.

Bozkurt, T. O., Schornack, S., Win, J., Shindo, T., Ilyas, M., Oliva, R., Cano, L. M., Jones, A. M. E., Huitema, E., & Kamoun, S. (2014). Rerouting of plant late endocytic trafficking toward a pathogen interface. Traffic, 15(9), 1027–1044.

Bozkurt, T. O., Schornack, S., Win, J., Shindo, T., Ilyas, M., Oliva, R., Cano, L. M., Jones, A. M. E., Huitema, E., van der Hoorn, R. A. L., & Kamoun, S. (2011). Phytophthora infestans effector AVRblb2 prevents secretion of a plant immune protease at the haustorial interface. Proceedings of the National Academy of Sciences of the United States of America, 108(51), 20832–20837.

Caillaud, M.-C., Wirthmueller, L., Sklenar, J., Findlay, K., Piquerez, S. J. M., Jones, J. D. G., & Faulkner, C. (2014). The plasmodesmal protein PDLP1 localises to haustoria-associated membranes during downy mildew infection and regulates callose deposition. PLoS Pathogens, 10, e1004496.

Camejo, D., Guzmán-Cedeño, Á., & Moreno, A. (2016). Reactive oxygen species, essential molecules, during plant-pathogen interactions. Plant Physiology and Biochemistry, 103, 10–23.

Chung IM, Venkidasamy B, Upadhyaya CP, Packiaraj G, Rajakumar G, Thiruvengadam M (2019) Alleviation of *Phytophthora infestans*-mediated necrotic stress in transgenic potato (*Solanum tuberosum* L.) with enhanced ascorbic acid accumulation. Plants 8:365

Cui, J., Luan, Y., Jiang, N., Bao, H., & Meng, J. (2017). Comparative transcriptome analysis between resistant and susceptible tomato allows the identification of lncRNA16397 conferring resistance to Phytophthora infestans by co-expressing glutaredoxin. Plant Biotechnology Journal, 89, 577–589.

D’Ambrosio, J. M., Couto, D., Fabro, G., Scuffi, D., Lamattina, L., Munnik, T., Andersson, M. X., Álvarez, M. E., Zipfel, C., & Laxalt, A. M. (2017). PLC2 regulates MAMP-triggered immunity by modulating ROS production in Arabidopsis. Plant Physiology, 174(4), 2363–2379.

Dagdas, Y. F., Belhaj, K., Maqbool, A., Chaparro-Garcia, A., Pandey, P., Petre, B., Tabassum, N., Cruz-Mireles, N., Hughes, R. K., Sklenar, J., Win, J., Menke, F., Findlay, K., Banfield, M. J., & Kamoun, S. (2016). An effector of the Irish potato famine pathogen antagonizes a host autophagy cargo receptor. eLife, 5, e10856.

Glazebrook, J. (2005). Contrasting mechanisms of defence against biotrophic and necrotrophic pathogens. Annual Review of Phytopathology, 43, 205–227.

Gonorazky, G., Guzzo, M. C., Abd-El-Haliem, A. M., Joosten, M. H. A. J., & Laxalt, A. M. (2016). Silencing of the tomato phosphatidylinositol-phospholipase C2 (SlPLC2) reduces plant susceptibility to Botrytis cinerea. Molecular Plant Pathology, 17(9), 1354–1363.

Guyon, A., Staps, T., Badot, L., & Schornack, S. (2025). Mutualist-pathogen co-colonisation modulates phosphoinositide signatures at host intracellular interfaces. bioRxiv.

Halim, V. A., Altmann, S., Ellinger, D., Eschen-Lippold, L., Miersch, O., Scheel, D., & Rosahl, S. (2009). PAMP-induced defense responses in potato require both salicylic acid and jasmonic acid. Plant Journal 57(2): 230–242. doi:10.1111/j.1365-313X.2008.03688.x

Heilmann, M., & Heilmann, I. (2022). Regulators regulated: different layers of control for plasma-membrane phosphoinositides in plants. Current Opinion in Plant Biology, 67, 102218.

Hirt, H. (1999). Transcriptional up-regulation of signalling pathways: more complex than anticipated? Trends in Plant Science, 4, 7–8.

Howe, G. A., Lightner, J., Browse, J., & Ryan, C. A. (1996). An octadecanoid pathway mutant (JL5) of tomato is compromised in signaling for defense against insect attack. The Plant Cell, 8(11), 2067–2077.

Juarez, M., et al. Effector Repertoire of a Dominant Phytophthora Infestans Isolate in Argentina and Its Relevance for Late Blight Resistance Deployment in Potato. Zenodo, 23 Sept. 2023

Judelson, H. S., & Ah-Fong, A. M. V. (2019). Exchanges at the plant-oomycete interface that influence disease. Plant Physiology, 179(4), 1198–1211.

Judelson, H. S., & Blanco, F. (2005). The spores of Phytophthora: weapons of the plant destroyer. Nature Reviews Microbiology, 3, 47–58.

Kadota, Y., Sklenar, J., Derbyshire, P., Stransfeld, L., et al. (2014). Direct regulation of the NADPH oxidase RBOHD by the PRR-associated kinase BIK1 during plant immunity. Molecular Cell, 54, 43–55.

Kamoun, S. (2006). A catalogue of the effector secretome of plant pathogenic oomycetes. Annual Review of Phytopathology, 44, 41–60.

Kiba, A., Nakano, M., Hosokawa, M., Galis, I., Nakatani, H., Shinya, T., Ohnishi, K., & Hikichi, Y. (2020). Phosphatidylinositol-phospholipase C2 regulates pattern-triggered immunity in Nicotiana benthamiana. Journal of Experimental Botany, 71(16), 5027–5038.

Kong, L., Ma, X., Zhang, C., Kim, S.-I., Li, B., Xie, Y., Yeo, I.-C., Thapa, H., Chen, S., Devarenne, T. P., Munnik, T., He, P., & Shan, L. (2024). Dual phosphorylation of DGK5-mediated PA burst regulates ROS in plant immunity. Cell, 187(3), 609–623.

Laxalt, A. M., Van Hooren, M., & Munnik, T. (2025). Phosphatidylinositol-specific phospholipase C in plant stress signalling: Emerging questions and new tools. Plant Physiology 197: 540–556.

Leesutthiphonchai, W., Vu, A. L., Ah-Fong, A. M. V., & Judelson, H. S. (2018). How Phytophthora infestans evades control efforts? Modern insight into the late blight disease. Phytopathology, 108(8), 916–924.

Li, L., Zhao, Y., McCaig, B. C., Wingerd, B. A., Wang, J., Whalon, M. E., Pichersky, E., & Howe, G. A. (2004). The tomato homolog of CORONATINE-INSENSITIVE1 is required for the maternal control of seed maturation, jasmonate-signaled defense responses, and glandular trichome development. The Plant Cell, 16(1), 126–143.

Liu, C., Zhang, Y., Tan, Y., Zhao, T., Xu, X., Yang, H., & Li, J. (2021). CRISPR/Cas9-mediated SlMYBS2 mutagenesis reduces tomato resistance to Phytophthora infestans. International Journal of Molecular Sciences, 22(21), 11423.

López-Gresa, M. P., Lisón, P., Yenush, L., Conejero, V., Rodrigo, I., & Bellés, J. M. (2016). Salicylic acid is involved in the basal resistance of tomato plants to Citrus Exocortis Viroid and Tomato Spotted Wilt Virus. PLOS ONE, 11(11), e0166938.

Luna, E., Pastor, V., Robert, J., Flors, V., Mauch-Mani, B., & Ton, J. (2011). Callose deposition: a multifaceted plant defence response. Molecular Plant-Microbe Interactions, 24, 183–193.

Mittler, R. (2017). ROS are good. Trends in Plant Science, 22, 11–19.

Mur, L. A. J., Kenton, P., Atzorn, R., Miersch, O., & Wasternack, C. (2006). Outcomes of concentration-specific interactions between salicylate and jasmonate signalling include synergy, antagonism and oxidative stress leading to cell death. Plant Physiology, 140, 249–262.

Otaiza-González SN, Mary VS, Arias SL, Bertrand L, Velez PA, Rodriguez MG, Rubinstein HR, Theumer MG (2022) Cell death induced by fumonisin B1 in two maize hybrids: correlation with oxidative status biomarkers and salicylic and jasmonic acids imbalances. Eur J Plant Pathol 163: 203–221.

Payá, C., Minguillón, S., Hernández, M., Miguel, S. M., Campos, L., Rodrigo, I., Bellés, J. M., López-Gresa, M. P., & Lisón, P. (2022). SlS5H silencing reveals specific pathogen-triggered salicylic acid metabolism in tomato. BMC Plant Biology, 22, 549.

Pavan, S., Jacobsen, E., Visser, R. G. F., & Bai, Y. (2010). Loss of susceptibility as a novel breeding strategy for durable and broad-spectrum resistance. Molecular Breeding, 25(1), 1–12.

Perk, E. A., Arruebarrena Di Palma, A., Colman, S., Mariani, O., Cerrudo, I., D’Ambrosio, J. M., Robuschi, L., Pombo, M. A., Rosli, H. G., Villareal, F., & Laxalt, A. M. (2023). CRISPR/Cas9-mediated phospholipase C2 knock-out tomato plants are more resistant to Botrytis cinerea. Planta, 257, 117.

Qi, F., Li, J., Ai, Y., Shangguan, K., Li, P., Lin, F., & Liang, Y. (2024). DGK5β-derived phosphatidic acid regulates ROS production in plant immunity by stabilizing NADPH oxidase. Cell Host & Microbe, 32(3), 425–440.

Qi, J., Wang, J., Gong, Z., & Zhou, J.-M. (2017). Apoplastic ROS signaling in plant immunity. Current Opinion in Plant Biology, 38, 92–100.

Qin, L., Zhou, Z., Li, Y., Li, M., Li, T., Zhang, L., Ding, X., & Luan, S. (2020). Specific recruitment of phosphoinositide species to the plant-pathogen interfacial membrane underlies Arabidopsis susceptibility to fungal infection. Plant Cell, 32(12), 3967–3988.

Rausche, J., Stenzel, I., Stauder, R., Fratini, M., Trujillo, M., Heilmann, I., & Rosahl, S. (2021). A phosphoinositide 5-phosphatase from Solanum tuberosum is activated by PAMP-treatment and may antagonize phosphatidylinositol 4,5-bisphosphate at Phytophthora infestans infection sites. New Phytologist, 229(1), 469–487.

Robert-Seilaniantz, A., Grant, M., & Jones, J. D. G. (2011). Hormone crosstalk in plant disease and defence: more than just jasmonate-salicylate antagonism. Annual Review of Phytopathology, 49, 317–343.

Robuschi, L., Mariani, O., Perk, E. A., Cerrudo, I., Villarreal, F., & Laxalt, A. M. (2024). Arabidopsis thaliana phosphoinositide-specific phospholipase C2 is required for Botrytis cinerea proliferation. Plant Science, 340, 111971.

Schornack, S., van Damme, M., Bozkurt, T. O., Cano, L. M., Smoker, M., Thines, M., Gaulin, E., Kamoun, S., & Huitema, E. (2010). Ancient class of translocated oomycete effectors targets the host nucleus. Proceedings of the National Academy of Sciences of the United States of America, 107(40), 17421–17426.

Schneider, C. A., Rasband, W. S., & Eliceiri, K. W. (2012). NIH Image to ImageJ: 25 years of image analysis. Nature Methods 9(7): 671–675.

Shimada, T. L., Betsuyaku, S., Inada, N., Ebine, K., Fujimoto, M., Uemura, T., Takano, Y., Fukuda, H., Nakano, A., & Ueda, T. (2019). Enrichment of phosphatidylinositol 4,5-bisphosphate in the extra-invasive hyphal membrane promotes Colletotrichum infection of Arabidopsis thaliana. Plant and Cell Physiology, 60(7), 1514–1524.

Simon, M. L. A., Platre, M. P., Assil, S., van Wijk, R., Chen, W. Y., Chory, J., Dreux, M., Munnik, T., & Jaillais, Y. (2014). A multi-colour/multi-affinity marker set to visualize phosphoinositide dynamics in Arabidopsis. The Plant Journal, 77(2), 322–337.

Smirnoff, N., & Arnaud, D. (2019). Hydrogen peroxide metabolism and functions in plants. New Phytologist, 221(3), 1197–1214.

Spoel, S. H., & Dong, X. (2008). Making sense of hormone crosstalk during plant immune responses. Cell Host & Microbe, 3, 348–351.

Thomazella, D. P. T., Brail, Q., Dahlbeck, D., & Staskawicz, B. J. (2021). CRISPR-Cas9-mediated mutagenesis of DMR6 orthologs in tomato confers broad-spectrum disease resistance. Proceedings of the National Academy of Sciences, 118, e2026152118.

Thordal-Christensen, H., Zhang, Z., Wei, Y., & Collinge, D. B. (1997). H₂O₂ accumulation in papillae and the hypersensitive response during the barley-powdery mildew interaction. Plant Journal 11(6): 1187–1194.

Torres, M. A., Jones, J. D. G., & Dangl, J. L. (2005). Pathogen-induced, NADPH oxidase-derived reactive oxygen intermediates suppress spread of cell death in Arabidopsis lesions. Plant Journal, 33, 299–316.

Turnbull, D., Yang, L., Naqvi, S., Breen, S., Welsh, L., Stephens, J., Morris, J., Boevink, P. C., Hedley, P. E., Zhan, J., Birch, P. R. J., & Gilroy, E. M. (2017). RXLR effector AVR2 up-regulates a brassinosteroid-responsive bHLH transcription factor to suppress immunity. Plant Physiology, 174(1), 356–369.

Van Schie, C. C. N., & Takken, F. L. W. (2014). Susceptibility genes 101: how to be a good host. Annual Review of Phytopathology, 52, 551–581.

Wang, Z., Su, C., Hu, W., Su, Q., & Luan, Y. (2023). The effectors of Phytophthora infestans impact host immunity upon regulation of antagonistic hormonal activities. Planta, 258, 59.

Weiß, L. S., Bradai, M., Bartram, C., Heilmann, M., Mergner, J., Kuster, B., Hensel, G., Kumlehn, J., Engelhardt, S., Heilmann, I., & Hückelhoven, R. (2024). Barley resistance and susceptibility to fungal cell entry involve the interplay of ROP signaling with phosphatidylinositol-monophosphates. bioRxiv.

Whisson, S. C., Boevink, P. C., Wang, S., & Birch, P. R. J. (2016). The cell biology of late blight disease. Current Opinion in Microbiology, 34, 127–135.

Yamamoto, Y. Y., Matsui, M., & Deng, X.-W. (1998). Positive feedback in plant signalling pathways. Trends in Plant Science, 3, 374–375.

Zhang, Y., Zhu, H., Zhang, Q., Li, M., Yan, M., Wang, R., Wang, L., Welti, R., Zhang, W., & Wang, X. (2009). Phospholipase Dα1 and phosphatidic acid regulate NADPH oxidase activity and production of reactive oxygen species in ABA-mediated stomatal closure in Arabidopsis. The Plant Cell, 21(8), 2357–2377.

Zhou, Z., Yu, D., Fan, B., Zhu, C., & Chen, Z. (2020). Phosphatidylserine synthase 1 is required for disease resistance and phosphatidylserine accumulation at the cell surface in Arabidopsis. New Phytologist, 226(2), 339–351.

