## Supplemental figures for "Phosphoinositide-specific Phospholipase C 2 (SlPLC2) Facilitates Vesicle Formation and Modulates Immune Signaling in Tomato *Phytophthora infestans* Interactions"

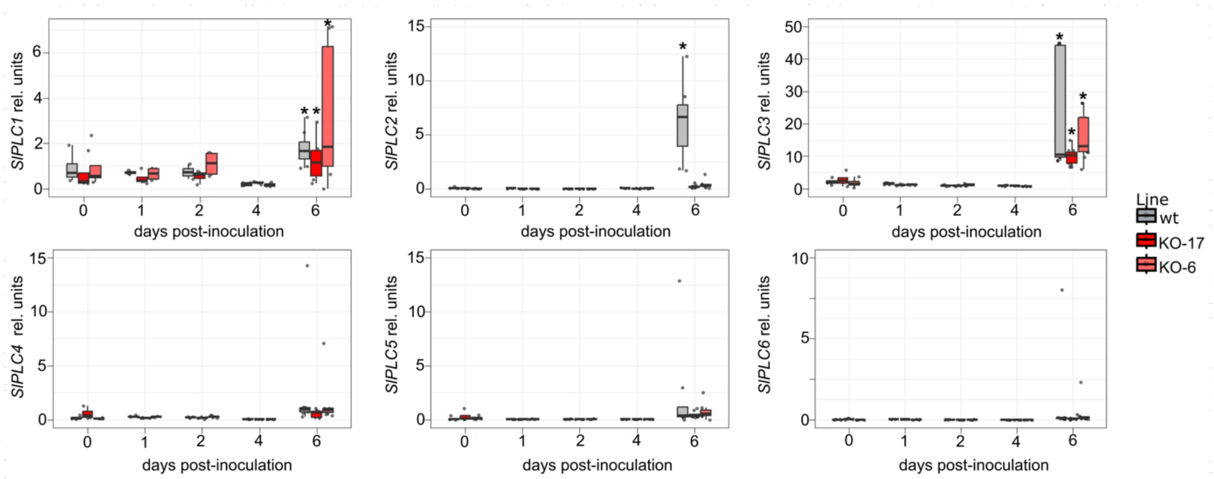


**Supplementary Figure S1. Expression of *SlPLC* gene family members during *P. infestans* infection.** Time-course RT-qPCR analysis of *SlPLC1–6* transcript levels in tomato leaflets inoculated with *P. infestans* (10 µl droplet of a 2x$10^5$ zoospore per mL). Samples were collected at 6 days post-inoculation (dpi) or from non-infected controls. Transcript levels were normalized to *SlACTIN* and expressed relative to non-infected samples. Data represent mean ± SE of four biological replicates (mock) and eight (infected). Asterisks indicate statistically significant differences compared to mock (Dunnett's test, *P* < 0.05).

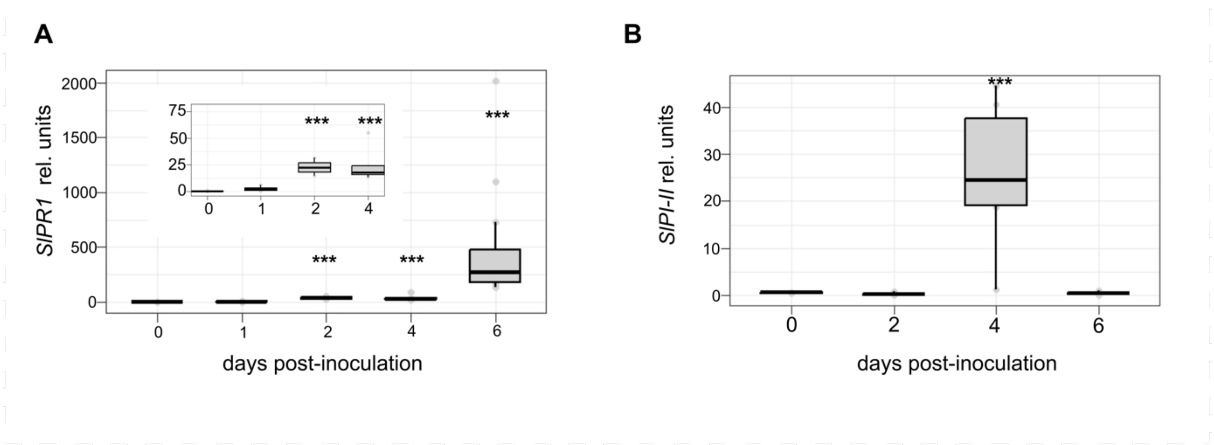


**Supplementary Figure S2. Salicylic acid and Jasmonic acid pathway marker expression during *P. infestans* infection.** **(A)** Time-course RT-qPCR analysis of *SlPR1a* transcript levels in wild-type (WT) tomato plants at 2, 4, and 6 dpi. **(B)** **(A)** Time-course expression of *SlPI-II* in WT plants at 2, 4, and 6 dpi, measured by RT-qPCR. Transcript levels were normalized to *SlACTIN*. Data represent mean ± SE of 4–12 biological replicates per time point.

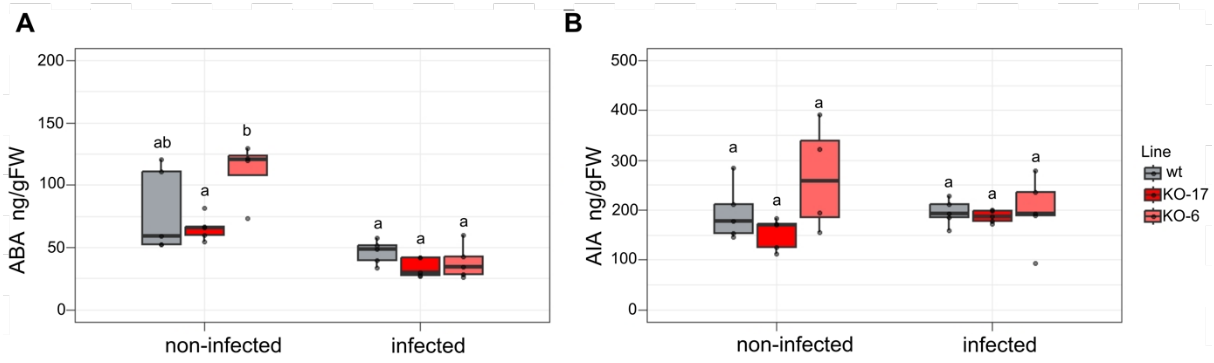


**Supplementary Figure S3. ABA and IAA remain unchanged in SlPLC2 knockout lines during *Phytophthora infestans* infection.** WT and SlPLC2 knockout tomato lines (KO-6 and KO-17) were inoculated with *P. infestans* zoospores by applying two 10 µL droplets of a 2 × $10^5$ zoospores/mL suspension on either side of the central vein of each leaflet. **(A–B)** Quantification of endogenous hormone levels by LC-MS/MS in aerial tissues at 6 dpi in infected or non-infected plants. Hormone concentrations were expressed as nanograms per milligram of fresh weight. **(A)** Abscisic acid (ABA), **(B)** Indole-3-acetic acid (IAA). Data represent the mean ± SE of biological replicates (n = 5). No statistically significant differences were detected between genotypes under either condition (Dunnett's test, P > 0.05).

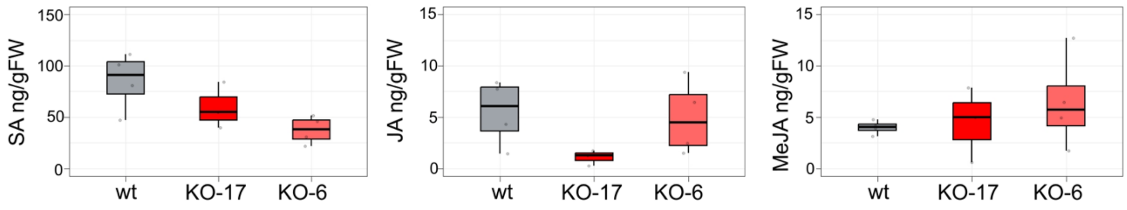


**Supplementary Figure S4. Basal levels of SA, JA, and MeJA are not altered in SlPLC2 knockout lines under non-infected conditions.** Detached leaflets from WT and SlPLC2 knockout tomato lines (KO-6 and KO-17) were processed immediately after excision, without any pathogen inoculation. **(A–C)** Quantification of endogenous hormone levels by LC-MS/MS in aerial tissues. Hormone concentrations were expressed as nanograms per milligram of fresh weight. **(A)** Salicylic acid (SA), **(B)** Jasmonic acid (JA), **(C)** Methyl jasmonate (MeJA). Data represent the mean ± SE of biological replicates (n = 5). No statistically significant differences were observed between genotypes (Dunnett's test, P > 0.05).

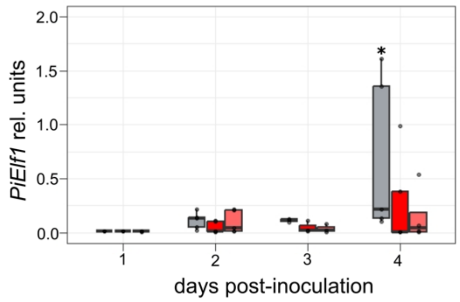


**Supplementary Figure S5. Quantification of *Phytophthora infestans* biomass in tomato leaflets over time.** Relative expression levels of the *P. infestans* elongation factor gene (*PiEF1-α*) were measured by RT-qPCR at 1, 2, 3, and 4 dpi in WT and KO tomato lines. Data were normalized to *SlACTIN* and expressed in arbitrary units. Box plots show median, interquartile range, and outliers (n ≥ 4). Asterisks indicate statistically significant differences compared to WT at each time point (ANOVA followed by Tukey's post-hoc test, $P < 0.05$).

45

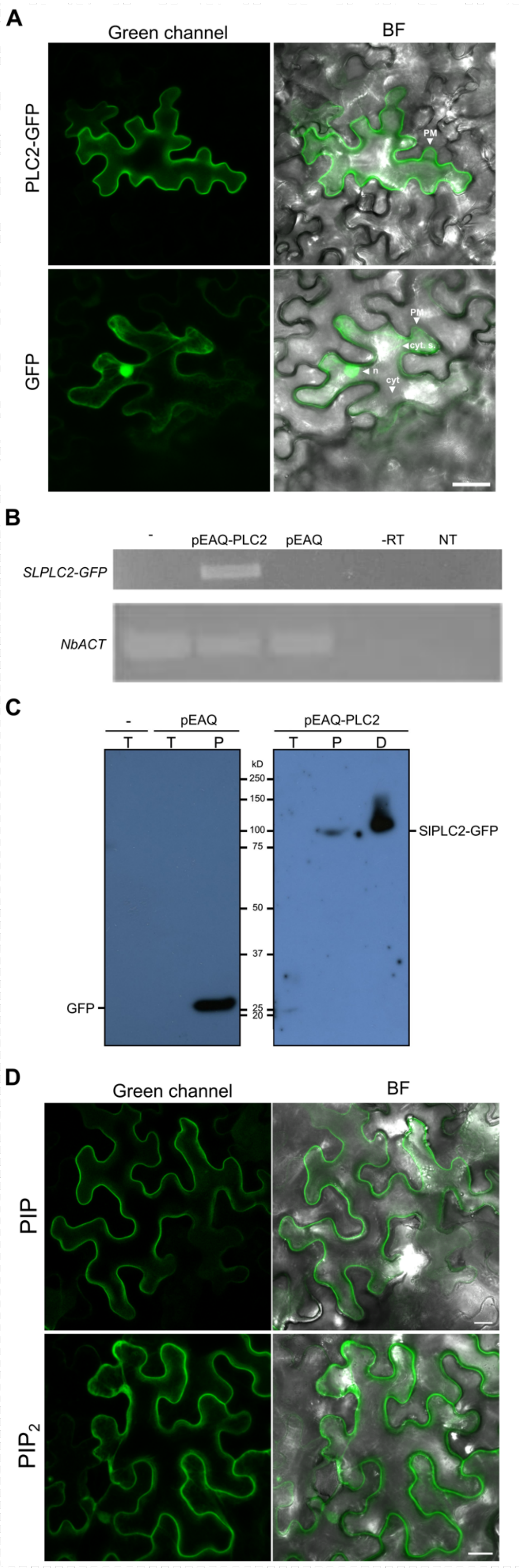
A
Green channel
BF
PLC2-GFP
PM
GFP
PM
cyt. s.
n
cyt
B
-
pEAQ-PLC2
pEAQ
-RT
NT
SLPLC2-GFP
NbACT
C
-
pEAQ
pEAQ-PLC2
T
T
P
T
P
D
kD
250
150
100
75
50
37
25
20
SlPLC2-GFP
GFP
D
Green channel
BF
PIP
PIP2

**Supplementary Figure S6. Localization and validation of SlPLC2-GFP expression in *Nicotiana benthamiana*, and distribution of lipid biosensors under basal conditions.** All experiments were conducted under non-infected (basal) conditions. **(A)** Confocal microscopy images showing subcellular localization of SlPLC2-GFP in epidermal cells. SlPLC2-GFP localizes predominantly to the plasma membrane. In contrast, free GFP displays diffuse cytosolic and nuclear localization. PM: plasma membrane; n: nucleus; cyt. s.: cytosolic strands; cyt: cytosol. Scale bar = 30 μm. **(B)** RT-PCR detection of the *SlPLC2*-GFP fusion transcript at 2 days post-infiltration confirms transcriptional expression. **(C)** Western blot analysis using anti-GFP antibody confirms protein-level expression of SlPLC2-GFP. Proteins were extracted from 4–5-week-old leaf tissue, separated on 10% SDS–PAGE, and transferred to nitrocellulose membranes. Ponceau S staining was used to verify equal loading. Membranes were incubated with alkaline phosphatase-conjugated anti-rabbit IgG secondary antibody. The positions of molecular weight markers (kDa) are indicated on the left. T: Total , P: Purified with His-tag column, D: Direct. **(D)** Confocal microscopy images showing the localization of PI4P and $PI(4,5)P_2$ biosensors in *N. benthamiana* epidermal cells. The PI4P biosensor (mCitrine-2xPH(FAPP1)) exhibits strong plasma membrane localization, while the $PI(4,5)P_2$ biosensor (mCitrine-2xPH(PLC)) displays both plasma membrane and cytosolic distribution. Images were acquired at 2 days post-infiltration. Scale bar = 20 μm.

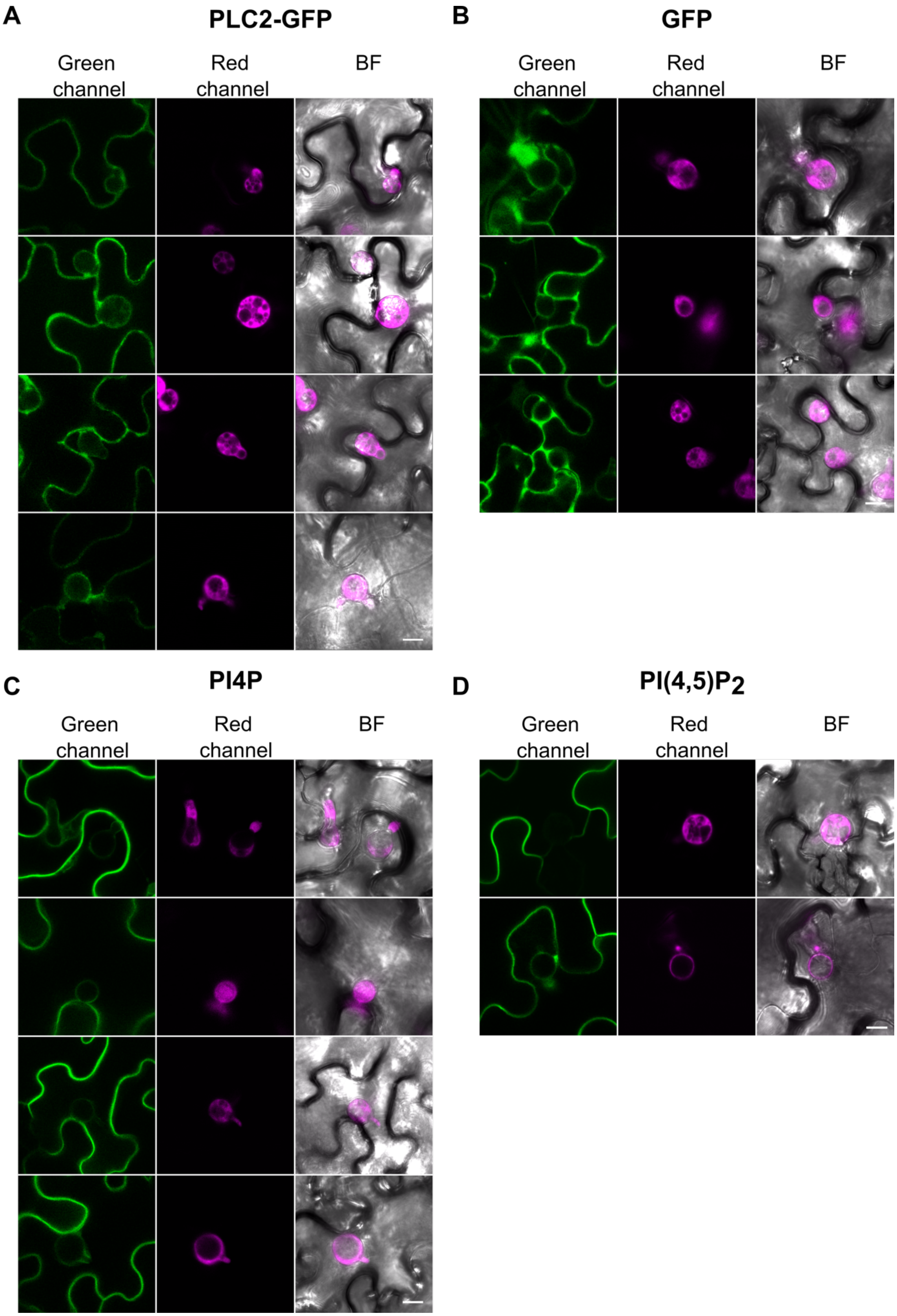
A
PLC2-GFP
Green channel
Red channel
BF
B
GFP
Green channel
Red channel
BF
C
PI4P
Green channel
Red channel
BF
D
PI(4,5)P2
Green channel
Red channel
BF

**Supplementary Figure S7. Subcellular localization of SlPLC2-GFP, free GFP, and phosphoinositide biosensors during *Phytophthora infestans* infection.** Confocal microscopy images of *Nicotiana benthamiana* epidermal cells at 2 days post-infiltration and 1-day post-inoculation with *P. infestans*. Leaf sectors were co-infiltrated with *Agrobacterium tumefaciens* carrying the indicated constructs and later inoculated with *P. infestans* zoospore suspension (2 × $10^5$ zoospores/mL; 10 µL per site). Whole-cell projections are shown to highlight subcellular distribution and vesicular localization. **(A)** SlPLC2-GFP: Plasma membrane signal with additional accumulation in infection-associated vesicular structures. **(B)** Free GFP: Diffuse cytosolic and nuclear signal, no specific enrichment in vesicles. **(C)** PI4P biosensor (mCitrine-2xPH(FAPP1)): Localizes to the plasma membrane and shows focal accumulations at infection sites. **(D)** PI(4,5)$P_2$ biosensor (mCitrine-2xPH(PLC)): Displays plasma membrane and cytosolic localization, with enriched labeling at the membranes surrounding infection vesicles. Scale bar = 20 µm.
